# Phenotypes of ultra-rare variant carriers benchmark variant effect scores

**DOI:** 10.64898/2026.10.01.755962

**Authors:** Shubhankar Londhe, Eva Holtkamp, Georgios Tsitsiridis, Anna Starovoit, Pedro Tomaz da Silva, Johannes Hingerl, Hilary Finucane, Julien Gagneur

## Abstract

Human genetics needs benchmarks for evaluating variant scores against observed phenotypic consequences. We therefore present UKBBGym, which evaluates coding and non-coding scores against plasma protein abundance and quantitative traits observed in carriers of ultra-rare variants in the UK Biobank. This design avoids ascertainment biases of clinical labels, potential physiological mismatch of experimental assays, and confounding pertaining to common-variant associations. UKBBGym recapitulates relative performance on pathogenicity prediction while enabling comparison across variant classes and molecular mechanisms. This shows that scores designed for coding and splicing variants correlate more strongly with protein abundance and quantitative traits than scores modelling transcriptional regulation. Moreover, experimental assays do not consistently outperform computational scores for predicting missense effects. Comparing captured to detectable variance indicates scope for improvement, particularly for indels and loss-of-function variants. Finally, for coding variants, UKBBGym can be reproduced from public summary statistics, providing an accessible population-phenotype benchmark complementary to clinical labels and experimental assays.

## Main

Identifying genetic variants that impair gene function is a central problem in genetics, with implications across many domains. In rare disease diagnostics, many disorders are caused by variants that reduce or abolish gene function. In oncology, somatic variants that impair tumor suppressor genes can drive tumorigenesis. Moreover, naturally occurring loss-of-function variants provide direct evidence linking gene impairment to phenotypic consequences, informing target validation and drug repurposing ^1^. Variant scores, which aim to quantify how strongly variants impair gene function, support variant prioritization in genetic diagnostics and oncology, and increase the power of statistical methods to identify genes involved in human traits ^2^.

However, predicting which variants impair gene function remains challenging. In rare disease diagnostics, the diagnostic rate typically remains below 50% across disorders ^3,4^. Improvement of in silico prediction could help reclassify variants and improve the diagnostic rate ^5^. Hence, variant scoring is an active field of research. A major class of variant scoring methods infers functional constraint from evolutionary conservation, ranging from alignment-based conservation scores (for instance, refs ^6,7^) to more recent genome- and protein-language models ^8–12^. Complementary approaches model specific biological processes, such as transcriptional regulation ^13–15^, splicing ^16–18^, RNA stability ^19^, or protein structure ^20,21^, and estimate the impact of variants by predicting disruption of these processes. A third class of variant scores integrates multiple sources of evidence (e.g., ^22,23^).

Critically, the evaluation and comparison of variant scores remain limited by the difficulty of establishing consistent and unbiased benchmarks. Existing benchmarks built on curated clinical variant databases such as ClinVar ^18^ and the Mendelian arm of TraitGym ^25^ provide variants with large, pathogenic effects. However, the extent of selection bias in these databases ^26^, particularly the underrepresentation of non-coding and hard-to-interpret variants, remains a limitation. Harmonization efforts, such as the ACMG/AMP variant interpretation guidelines ^27,28^, improve classification consistency but do not remove this selection bias. Complementary to clinical variant databases, multiplexed assays of variant effects provide dense functional measurements under controlled experimental conditions ^29,30^. However, harmonizing heterogeneity across assays and their readouts is challenging and remains an ongoing effort ^31,32^. Furthermore, these in vitro measurements may not fully capture phenotypic consequences observed in carriers of these variants. Variant effects in humans can be assessed through associations with common variants, as in eQTL- and GWAS-based benchmarks ^13,25^. However, fitness effects of common variants are small ^33^, and the evaluation of these effects is subject to confounding due to linkage disequilibrium, shared genetic structure, and environmental factors, among others ^34^. Rare variants alleviate these problems and have been used directly to rank variant scores by their correspondence with traits of variant carriers in large cohorts, but this approach has only been explored for missense variants ^35,36^.

To address these limitations, we present UK Biobank Gym (UKBBGym), a benchmark that evaluates variant scores against phenotypes observed in human carriers of ultra-rare variants from the whole-genome sequenced UK Biobank cohort ^37,38^. By leveraging systematically collected phenotypes from a population-based cohort, UKBBGym reduces the ascertainment and reporting biases inherent in clinically curated variant and disease datasets. We focus on quantitative phenotypes rather than binary disease phenotypes, which allows us to leverage the continuum of variant effects across a population. Moreover, by focusing on ultra-rare variants, UKBBGym captures variants with large phenotypic effects and mitigates spurious correlations pertaining to common variant associations. UKBBGym evaluates variant scoring methods against readouts from two complementary phenotypic modalities: i) the protein abundances of 1,076 genes from Olink plasma proteomics ^39^ and ii) quantitative traits, including blood biochemistry, anthropometry, and hematology for 670 gene-trait pairs. These modalities allow us to compare variant scores against gene expression and quantitative traits. For usability, the benchmarking workflow is implemented in two versions. The primary workflow is implemented on the UK Biobank Research Analysis Platform for approved users. A version restricted to quantitative traits using summary statistics from the Genebass study ^40^ allows a broader set of users to benchmark methods on coding variants.

## Results

### Population-scale phenotypes benchmark variant scores against plasma protein abundance and quantitative traits

UKBBGym benchmarks variant scores against two population-scale phenotypic readout modalities: plasma protein abundance and quantitative traits. We considered ultra-rare variants (allele count ≤ 20) within ±5 kb of the MANE transcript in the UK Biobank whole-genome sequencing ^38^. This yields a variant set comparable in size to ClinVar in coding regions but orders of magnitude larger in enhancers, promoters, and introns, where existing benchmarks are sparse (Fig. 1a).

**Figure 1.**
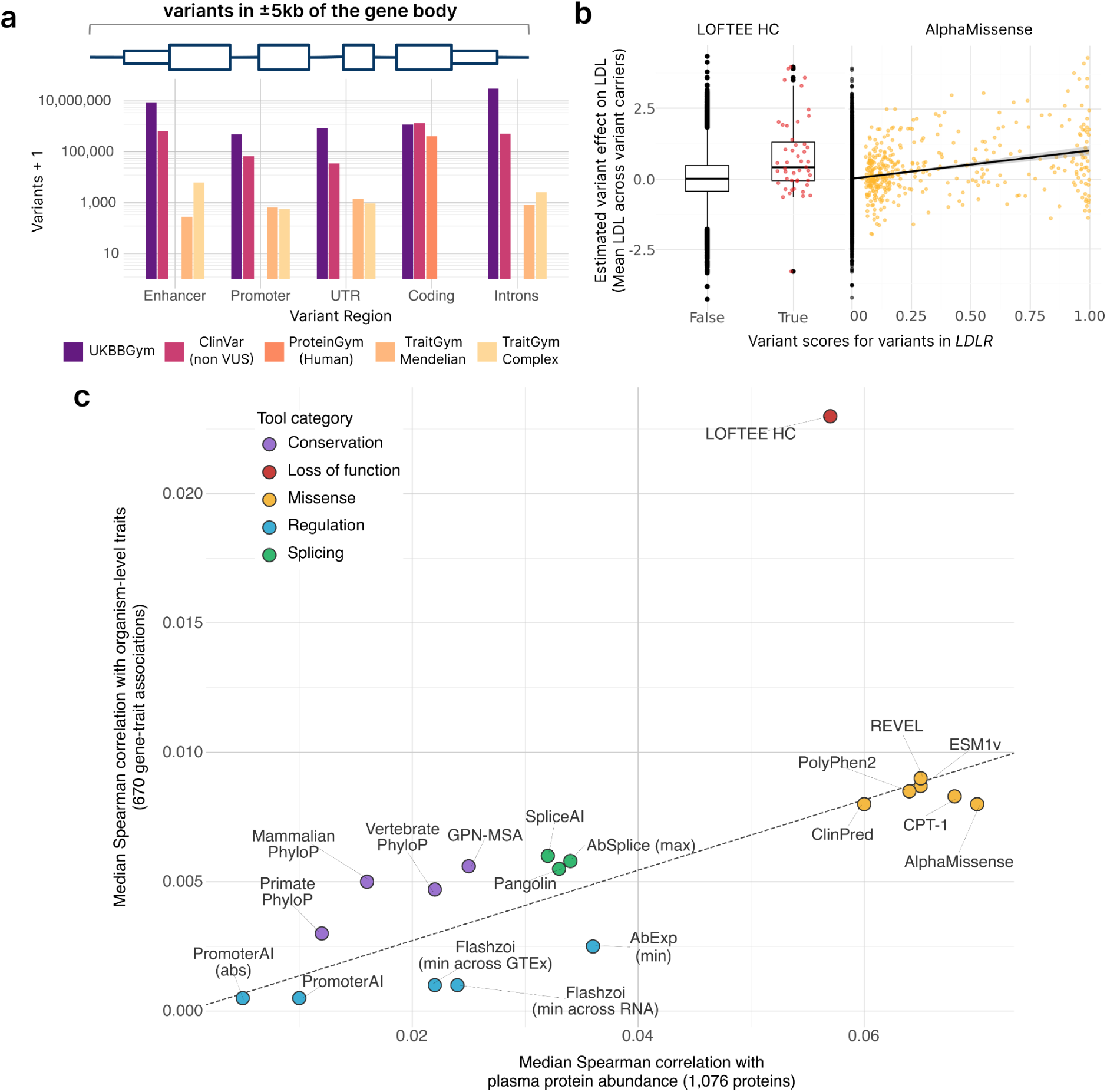
Population-scale phenotypes benchmark variant scores against plasma protein abundance and quantitative traits. **a**, Top: Schematic of the variant selection and gene-region definition. We considered ultra-rare variants (allele count ≤ 20) within ±5kb of the MANE transcript of coding genes and the following gene regions: ENCODE enhancers, promoters (TSS ± 500 bp), UTR, coding, and introns. Bottom: Number of variants used in variant effect prediction benchmarks per genomic region for UKBBGym (purple), ClinVar (excluding variants of uncertain significance, burgundy), ProteinGym human (orange), TraitGym Mendelian (peach), and TraitGym Complex (yellow). **b,** LDL cholesterol variant effect (y-axis), defined as the mean of adjusted LDL cholesterol level across variant carriers, against variant scores (x-axis) using LOFTEE high confidence (boxplot, left) and AlphaMissense (scatterplot, right) for *LDLR* variants. For AlphaMissense, variants without a missense score were assigned the neutral default score (Methods). The line shows a linear regression fit for visualization. For the LOFTEE boxplot, the center line indicates the median; box limits indicate the first and third quartiles; whiskers span all data within 1.5 interquartile ranges of the lower and upper quartiles. **c,** Median per-gene Spearman correlation between variant score and variant effect estimates on quantitative traits across 670 gene-trait pairs (y-axis) against median per-gene Spearman correlation between variant scores and variant effect estimates on plasma protein abundance across 1,076 genes (x-axis). For comparability, this analysis is restricted to single-nucleotide variants. Each point represents one variant scoring method and is colored by functional category. The dashed line indicates the median ratio of trait-to-protein-abundance correlations across scoring methods. Loss-of-function annotations were used to define the gene-trait pairs, which may have introduced circularity in the correlation with LOFTEE high confidence.

For the quantitative traits, we identified gene-trait pairs by burden testing LOFTEE high confidence (HC) loss-of-function variants ^41^ against biochemical, anthropometric, and hematological quantitative traits adjusted for age, sex, genetic principal components, and common-variant polygenic risk ^42^. We performed association testing using ultra-rare variants in 379,645 unrelated individuals of European ancestry to limit confounding associations from population structure and relatedness. We restricted the analysis to single-nucleotide variants (SNVs) and indel variants of less than 50 bp that lie within ±5 kb of the MANE transcript. For each gene, we retained the most strongly correlated trait to prevent the benchmark from being biased toward genes with many associated traits. In total, we obtained 670 gene-trait pairs spanning 96 quantitative traits (false discovery rate < 0.05; Supplementary Table 1). Furthermore, we adjusted Olink plasma protein abundance ^39^ measurements for technical and biological variation using the autoencoder-based PROTRIDER ^43^. Among the methods tested, PROTRIDER was best enriched for ultra-rare variants likely to affect protein abundance. Specifically, 15.5% (95% CI [13.4, 17.6]) harbored a rare LOFTEE HC variant among the 1,000 top-ranked adjusted values, compared to 5.6% (95% CI [4.2, 6.9]) for covariate regression residuals (Supplementary Fig. 1). Following Parry et al. ^36^, we retained genes in which loss-of-function variants significantly reduced adjusted protein abundance, yielding 1,086 genes, of which 1,076 had at least 100 variants (Supplementary Table 2).

Having identified bona fide gene-trait pairs and gene-protein abundance pairs, we established the benchmarking metrics. We considered the same set of variants as for the association testing. We further restricted to heterozygous alleles to not confuse the analysis with dominance or recessive effects, which are outside the scope of the vast majority of variant-scoring methods.

For each variant, we estimated the variant effect as the average phenotype (be a quantitative trait or protein abundance) across its carriers and rank-correlated it with various scores, both qualitative and quantitative (Supplementary Table 3). Because phenotypes were centered and the variants were ultra-rare, the mean phenotype among carriers is effectively equal to the difference between carriers and non-carriers. Illustrative of our benchmark, variants annotated as high-confidence loss-of-function by LOFTEE HC in the low-density lipoprotein receptor gene (*LDLR*) associated with elevated low-density lipoprotein (LDL) levels in blood, compared to non-LOFTEE HC variants (Spearman ρ = 0.037; Fig. 1b). In comparison, scoring variants with the AlphaMissense ^20^ pathogenicity continuous score, whereby AlphaMissense scores were set to 0 for non-missense variants, yielded a stronger rank-correlation (ρ = 0.066; Fig. 1b). Like AlphaMissense, many methods do not annotate all possible variants. For these unannotated variants, we assigned a default score equal to the method’s neutral effect score (Methods). This allows comparison of methods across variant categories. To align the direction of effects across genes, we reversed the phenotype sign where necessary so that LOFTEE HC loss-of-function variants were associated with positive adjusted phenotypic values. Rank correlation was also used for protein abundances. To improve robustness, we restricted our analyses throughout the study to correlations computed across at least 100 variants.

By benchmarking loss-of-function, missense, splicing, regulatory, and conservation-based predictors on a common scale, UKBBGym reveals that predictor classes differ not only in overall performance but also in whether they relatively preferentially capture effects on plasma protein abundance or quantitative traits. This comparison was restricted to SNVs, which all scoring methods can annotate (Fig. 1c, pairwise statistical comparisons in Supplementary Fig. 2). Specifically, loss-of-function annotation showed the strongest correlation with quantitative traits. However, this observation is partly circular because loss-of-function annotation was used to define gene-phenotype pairs and to orient effect directions. Missense predictors, including CPT-1 ^44^, AlphaMissense ^20^, ESM1v ^45^, and REVEL ^46^, showed the strongest correlations with plasma protein abundance and strong correlations with traits. Splicing predictors formed an intermediate group, with SpliceAI ^16^, Pangolin ^17^, and AbSplice ^18^ showing moderate correlations with protein abundance and intermediate correlations with traits. Conservation-based scores displayed a distinct profile, correlating only modestly with protein abundance but comparatively well with quantitative traits. This suggests that evolutionary constraint captures dimensions of functional impact that are not fully reflected in plasma protein levels. The two RNA expression models, PromoterAI ^14^ and Flashzoi ^13,47^, and to a lesser extent the aberrant expression model AbExp ^48^, showed modest correlations with protein abundance but little or no correlation with traits. Because PromoterAI and Flashzoi were applied only to SNVs in predefined promoters and enhancers owing to their computational demands, their overall performance and utility may be underestimated.

Collectively, the biological class of variant effect being modeled was a stronger determinant of performance than the choice of individual predictor within a class. Moreover, variant scores have distinct yet correlated predictive power for plasma protein abundance and quantitative traits, and methods that score coding and splicing variants correlate more strongly with both readouts than those that score transcriptional regulation.

### Missense variant score performance depends on the training strategy and the protein region

We next examined missense predictors in greater detail, asking how their performance depends on training strategy and the structural context of the affected residue.

Some missense predictors are trained on clinically reported variants to discriminate pathogenic from benign variants, a strategy that has raised concerns of bias ^32,49^. As representative tools of this category, we considered ClinPred ^50^, REVEL, PolyPhen2 ^51^, and BayesDel ^52^. We contrasted those tools against missense predictors that are not trained on clinical labels: AlphaMissense, CPT-1, and ESM1v. Furthermore, we considered general conservation scores: Vertebrate PhyloP ^6^ and GPN-MSA ^11^. Finally, we considered the integrative deleteriousness score CADD ^22,53^, which combines diverse genomic annotations and is trained to discriminate between variants fixed in human populations and simulated de novo variants.

On the protein abundance track, CPT-1 achieved the highest Spearman correlation, closely followed by AlphaMissense and ESM1v (Fig. 2a). The same overall hierarchy was preserved on quantitative traits across 581 gene-trait pairs (across genes with at least 100 missense variants), with CPT-1 again leading (Fig. 2b). Clinical-label-trained tools showed lower Spearman correlations than non-clinically-trained predictors. This gap between clinical and non-clinical-label-trained methods, evident in both the Spearman correlation and the mean variant effects among top-ranked variants, is consistent with suspected biases in clinically trained predictors ^32,49,54^. Conservation scores (GPN-MSA, Vertebrate PhyloP) along with PolyPhen2 showed the lowest correlations with both quantitative traits and protein abundances.

**Figure 2.**
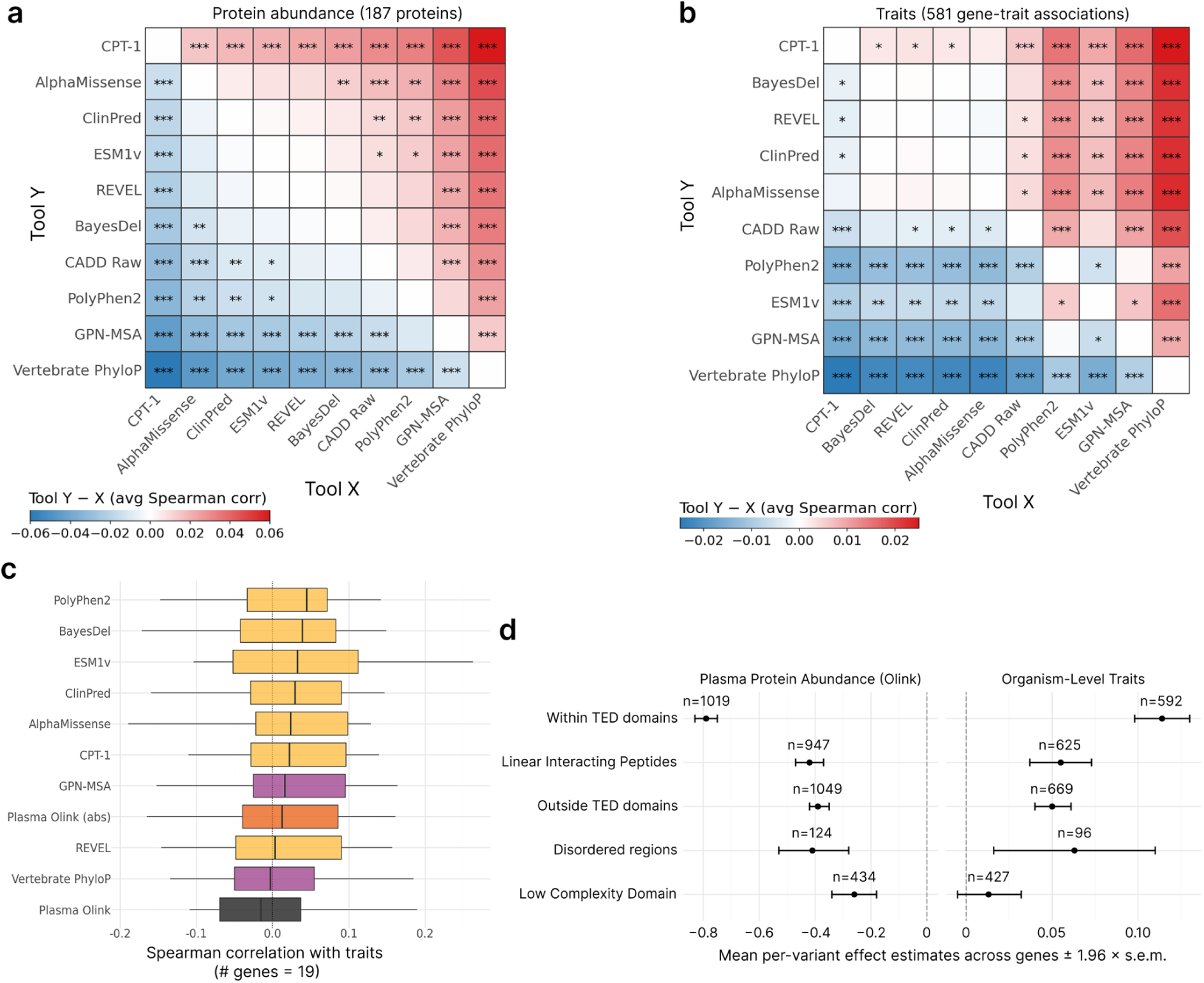
Missense variant score performance depends on training strategy and protein region. **a**, Pairwise comparison of missense variant scores based on their per-protein Spearman correlations with plasma protein abundance across the genes with at least 100 ultra-rare missense variants (*n* = 187). Each cell shows the mean paired difference in the Spearman correlation between the row and column scores. Red indicates the row score correlates more strongly with abundance than the column score, and blue indicates the reverse. Significance of the paired correlations is assessed by the Wilcoxon signed-rank test across proteins (*** *P* < 0.001, ** *P* < 0.01, * *P* < 0.05). **b,** As in (a) for quantitative traits across gene-trait pairs (*n* = 581). **c,** Distribution of the Spearman correlations between quantitative traits and missense variant scores. The mean of adjusted protein abundance across variant carriers (black) and its absolute value (orange) are considered as variant scores (*n =* 19 gene-trait pairs with at least 100 ultra-rare variants and plasma protein abundance measurements). The centerline indicates the median; box limits indicate the first and third quartiles; whiskers span all data within 1.5 times the interquartile range of the lower and upper quartiles. **d,** Mean across genes of estimated variant effects (left: protein abundance, right: traits) averaged within each gene, stratified by protein region: within TED domains, outside TED domains, disordered regions (from MobiDB), linear interacting peptides (from MobiDB), and low-complexity domains (LCD-composer). Error bars, 1.96 × s.e.m. across genes.

Given that variant scores correlated with both protein abundance and quantitative traits, we next asked whether measured plasma protein abundance could substitute for genotype-based deleteriousness predictions when ranking estimated effects of ultra-rare missense variants on quantitative traits. Across 19 gene-trait pairs with at least 100 variants in carriers with the corresponding protein abundance measured in plasma, per-variant protein abundance showed weak rank correlation with quantitative-trait variant effect estimates and was outperformed by the missense deleteriousness predictors (Fig. 2c). This observation is consistent with studies showing that the effects of molecular intermediates, such as gene expression on downstream quantitative phenotypes, are often attenuated or incompletely predictive ^55,56^. Nonetheless, the variants with the largest abundance deviations were associated with quantitative-trait variant effects comparable to those prioritized by the best computational scores (Supplementary Fig. 3). Thus, among individuals carrying ultra-rare missense variants in these genes, variant scores tracked quantitative traits more closely than measured protein abundance, although the most extreme plasma protein abundance levels remained informative.

Moreover, variants in structured domains annotated by The Encyclopedia of Domains ^57^ (TED) were associated with the largest average variant effect estimates on plasma protein abundance and quantitative traits, whereas variants in disordered, low-complexity, and other non-structured regions generally were associated with milder phenotypes (annotated by MobiDB ^58^ and LCD-Composer ^59^, respectively; Fig. 2d).

Altogether, these results show that UKBBGym separates missense predictors by training strategy, that measured plasma protein abundance does not substitute for genotype-based prediction, and that the associated effects depend on the structural context of the affected residue.

### Scores for coding and splice-altering variants capture the strongest protein-abundance effects

UKBBGym provides an opportunity to study the effects of coding and non-coding variation on gene expression at the level of protein abundance, and to benchmark variant scores against these effects. ^55,56^.

We first compared how protein abundance associated with non-coding SNVs and indels. Although SNVs accounted for most rare non-coding variation, their associations with plasma protein abundance were generally modest. By contrast, indels were associated with substantially larger abundance reductions across size classes and regulatory contexts (Fig. 3a). For scoring indels, current sequence-based models of gene expression require adaptations and remain computationally impractical at the biobank scale ^60^, leaving this high-impact class of non-coding variation poorly covered by existing predictors in practice.

**Figure 3.**
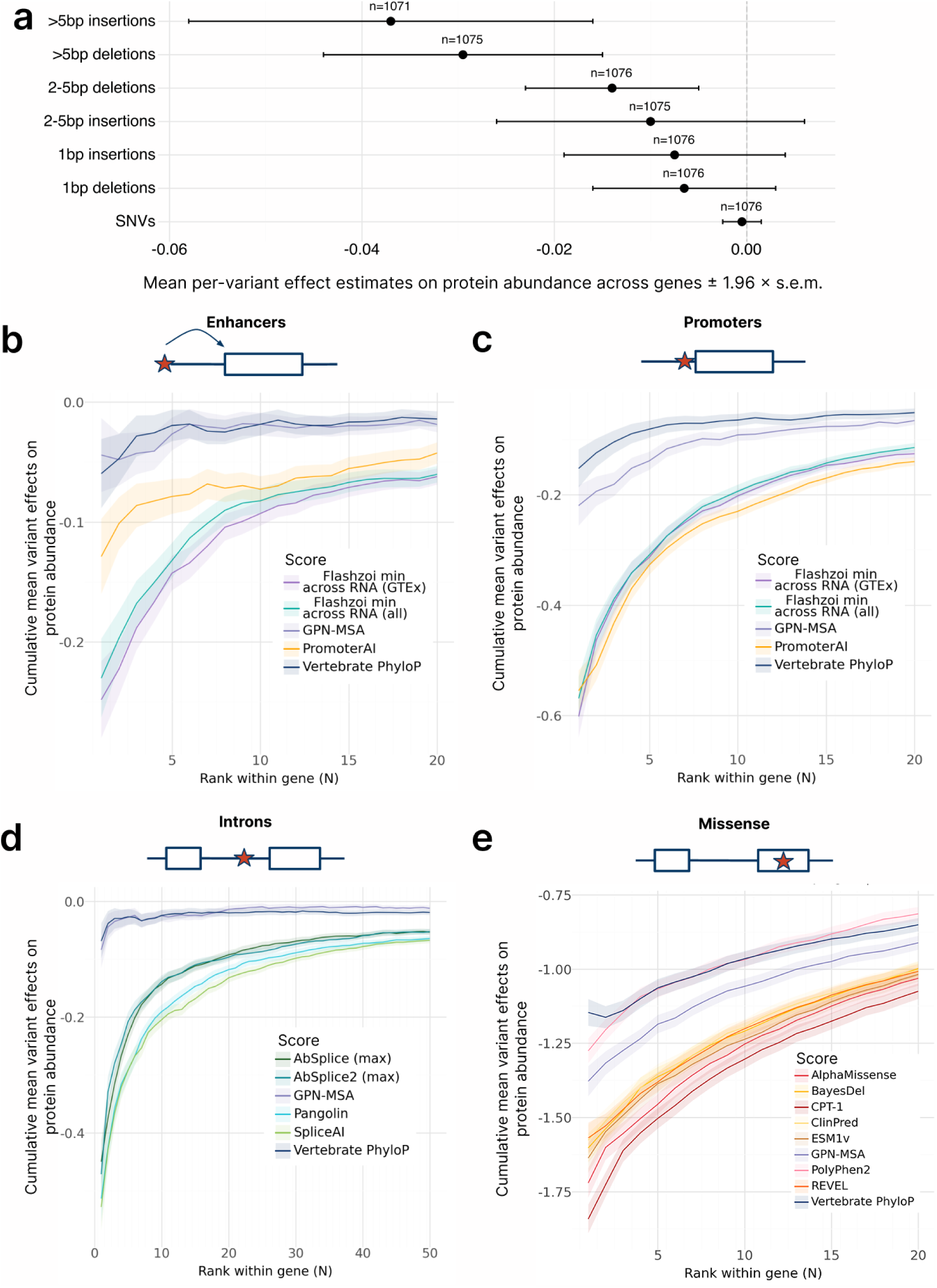
Scores for coding and splice-altering variants capture the strongest protein-abundance effects. **a**, Mean across genes of estimated variant effects on protein abundance averaged within each gene, stratified by non-coding variant length. Sample sizes indicate the number of genes; error bars represent 1.96 × s.e.m. across genes. **b,** Cumulative mean variant effect on protein abundance (y-axis) across the top-ranked variants (x-axis) according to various scores for ENCODE-annotated enhancer variants (pELS and dELS). Ribbons indicate 1× s.e.m. across genes. **c,** Same as in (b) but restricted to promoter variants (TSS ± 500 bp). **d,** Same as in (b) but restricted to intronic variants, excluding splice site dinucleotides. **e,** Same as in (b) but restricted to missense variants.

We therefore focused on scores for SNVs, excluding enhancer and promoter variants that overlapped the coding sequence or were predicted to affect splicing (Methods). Because Spearman correlation summarizes ranking performance across the full spectrum of variants, we complemented it with a top-ranked variant analysis that shows the mean estimated effect across the most highly prioritized variants by each method in each gene. This provides an interpretable view of each score’s ability to identify the most consequential variants within a gene. The relative ordering of scores was generally stable across rank thresholds, indicating that the comparison was not driven by an arbitrary choice of cutoff. Within ENCODE-annotated enhancers ^61^, Flashzoi outperformed PromoterAI, the conservation scores GPN-MSA, and vertebrate PhyloP in prioritizing SNVs associated with reduced protein abundance (Fig. 3b). PromoterAI was included in this analysis because its input regions overlapped many of these enhancers. Moreover, Flashzoi and PromoterAI demonstrated comparable performance in PromoterAI’s promoter regions (TSS ± 500 bp), where both outperformed conservation-based scores (Fig. 3c).

We next turned to intronic SNVs, excluding promoter, enhancer, and core splice-site dinucleotides, and evaluated splicing predictors designed to identify variants that alter splicing, a major mechanism through which intronic variation can affect RNA and protein abundance. The splicing-specific tools SpliceAI and Pangolin, followed by AbSplice (using the maximum predicted effect across GTEx tissues), consistently outperformed conservation scores in predicting plasma protein abundance (Fig. 3d).

Considering missense SNVs, CPT-1 prioritized the variants with the largest associated reductions in plasma protein abundance, followed by AlphaMissense and ESM1v. Apart from PolyPhen2, the missense-specific predictors ranked variants with lower protein abundance than the conservation scores GPN-MSA and Vertebrate PhyloP (Fig. 3e). Clinical-label-trained (ClinPred, BayesDel, REVEL) tools showed lower Spearman correlations than non-clinically-trained predictors and, at the top of their rankings, prioritized variants associated with less extreme effects on both protein abundance and quantitative traits (Supplementary Fig. 4).

Strikingly, the protein abundance reductions associated with top-ranked SNVs differed markedly across contexts, reaching approximately 0.2 standard deviations in enhancers, 0.6 in promoters, and 0.5 in introns (Fig. 3b-d). Mean estimated variant effect on protein abundance likewise decreased with distance from the transcription start site, from core promoter and promoter regions to TF-binding and enhancer regions, where mean effects were near zero (Supplementary Fig. 5). Part of the difference between contexts therefore reflects the underlying variant effects rather than predictor performance alone. In contrast to intronic SNVs ranked by splicing predictors, which are also associated with traits, promoter and enhancer SNVs prioritized by Flashzoi and PromoterAI did not show significant effects on traits (Supplementary Fig. 6), consistent with the low Spearman correlations with quantitative traits reported above.

Altogether, these comparative analyses established a hierarchy of variant categories and scores associated with extreme reductions in protein abundance. Indels, particularly larger indels, showed greater reductions than SNVs, while among SNVs, scores targeting coding and splice-altering variation identified more extreme abundance reductions than models of transcriptional regulation in promoters, and especially enhancers.

### UKBBGym is reproducible from summary statistics and reveals variant-class-specific prediction gaps

Access to individual-level biobank data can limit the usability of phenotype-based benchmarks. Prompted by the temporary, community-wide suspension of access to the UK Biobank Research Analysis Platform in April 2026, we asked whether UKBBGym could, in part, be reconstructed using published single-variant association test statistics from 394,841 UK Biobank exome sequencing data (Genebass ^40^). For ultra-rare variants, the additive effect from a standard association model approximates the average phenotype across variant carriers up to a scaling factor that is negligible at the allele counts considered (Methods). This equivalence allowed the analyses for coding variants to be reproduced without individual-level data.

Consistently, pairwise rankings of missense scoring methods obtained from Genebass effect estimates (Fig. 4a) agreed with those obtained from individual-level data. To expand our initial analysis, we added eleven variant scores to the missense analyses: the recently released AlphaGenome Variant Impact (AVI) ^23^, GPN-Star Vertebrate (V), Mammalian (M), and Primate (P) ^12^, as well as popEVE ^62^, MSA Pairformer ^9^, SIFT ^63^, Mammalian and Primate PhyloP ^6^, and gnomAD exome and genome allele frequencies ^64,65^. We also added protein region annotations covering structured domains, interaction interfaces, and disordered and low-complexity regions. Across 475 gene-trait pairs in genes with more than 100 missense variants, CPT-1, AVI, popEVE, and AlphaMissense correlated most strongly with Genebass effect estimates, and were not distinguishable from one another. All four were significantly above the conservation-based variant scores and the gnomAD allele frequency, which ranked lowest.

**Figure 4.**
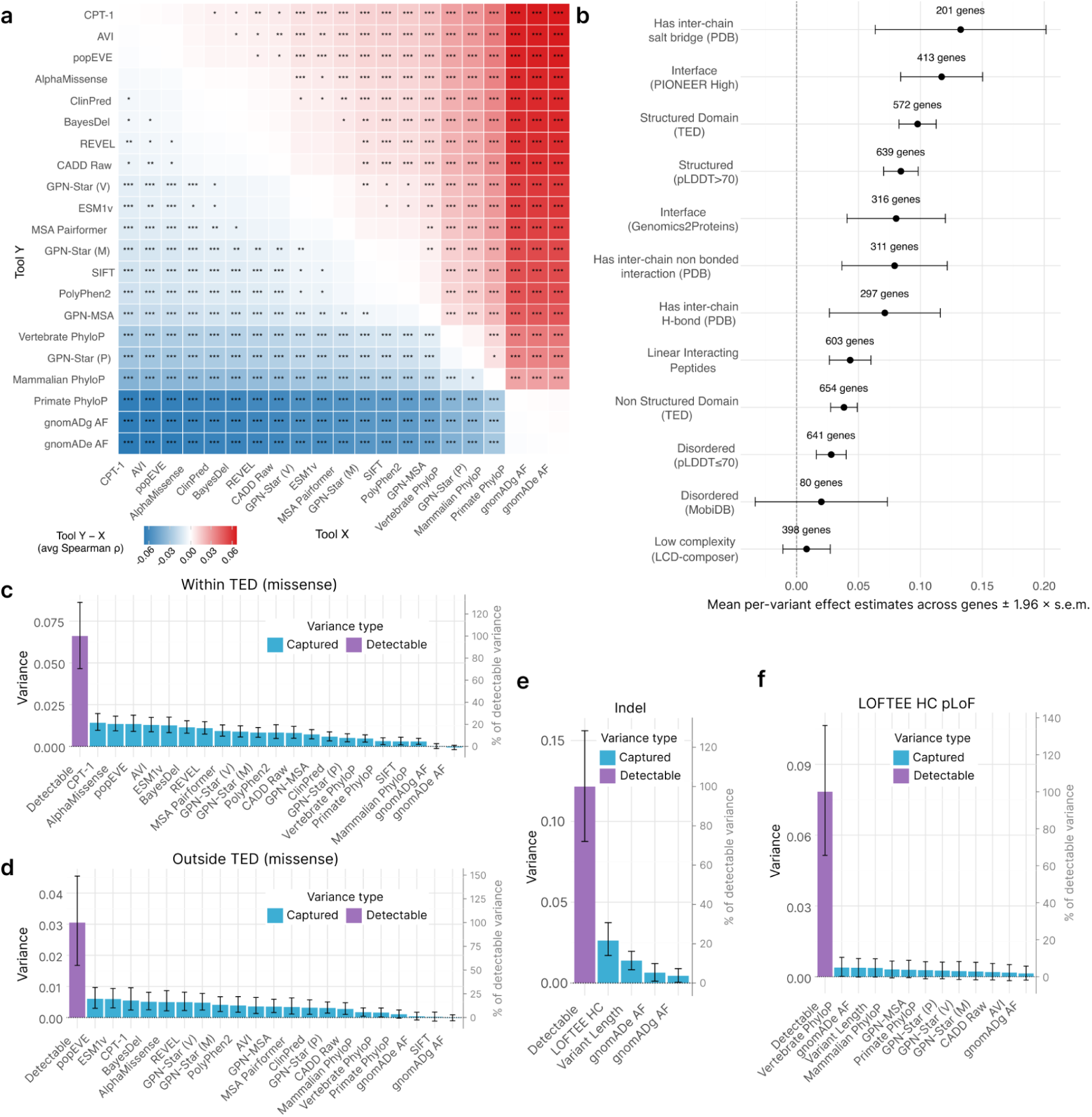
UKBBGym is reproducible from summary statistics and reveals variant-class-specific prediction gaps. **a**, Pairwise comparison of missense variant scoring methods based on their Spearman correlations with Genebass variant effect estimates for variants in 475 gene-trait pairs. Each cell compares the row method (Tool Y) against the column method (Tool X): color shows the mean paired difference in Spearman correlation (Tool Y − Tool X). Red indicates the row method correlates more strongly with effect estimates than the column method, and blue indicates the reverse. Significance of the paired correlations is assessed by the Wilcoxon signed-rank test across genes (*** *P* < 0.001, ** *P* < 0.01, * *P* < 0.05). **b,** Mean across genes of Genebass variant effect estimates averaged within each gene, for missense variants stratified by protein structural and interaction annotation. Error bars indicate 1.96 × s.e.m. **c,** Variance in variant effect estimates captured by each variant score (primary y-axis) for variants within TED-annotated domains. The purple bar indicates the detectable variance, defined as the observed variance minus the estimated sampling-error variance. The percentage of the detectable variance is shown on the secondary y-axis. Error bars indicate 95% confidence intervals from 2,000 gene-level bootstraps. **d,** Same as in (c) but for variants outside TED-annotated domains. **e,** Same as in (c) but for insertion and deletion (indel) variants. **f,** Same as in (c) but for putative loss-of-function variants classified as LOFTEE HC.

We also considered the All by All summary statistics from 392,030 participants from the All of Us cohort ^66^ and found no contradictions between pairwise comparisons of variant scores with the Genebass-based analysis (Supplementary Fig. 7). However, All by All had fewer trait measurements and lower per-trait coverage than Genebass, resulting in fewer gene-trait pairs (260 vs. 670, Supplementary Table 4) and therefore less power to distinguish between methods. We therefore focused the subsequent analyses on Genebass and, consequently, on coding variants and quantitative traits.

Clinically curated pathogenic variants have been reported to be enriched at interaction interfaces ^67,68^ and depleted in disordered regions, where predictors show lower sensitivity, potentially inflating apparent performance when ordered and disordered regions are evaluated together ^69^. In contrast, population-phenotype associations are less prone to this form of ascertainment bias because phenotype collection does not directly depend on prior knowledge or variant classification. We found that the effect estimates were highest for variants at inter-chain salt bridges and high-confidence PIONEER interfaces ^68^ (Fig. 4b). Variants in structured regions followed, whether defined as TED-annotated domains (Supplementary Fig. 8a, b) or by AlphaFold2 ^70^ pLDDT>70 (Supplementary Fig. 8c, d). Variants in disordered regions showed lower effect estimates, and variants in low-complexity regions the lowest, with intervals overlapping zero. The same ordering held for plasma protein abundance. Hence, these findings confirm in population phenotypes that missense variants in ordered protein regions are associated with larger effects than those in disordered regions, and quantify the magnitude of this difference.

We next asked how much of the overall variation that is in principle distinguishable from sampling error is captured by current variant scores. Because phenotypic measurements from multiple individuals carrying the same rare allele may not be statistically independent, we restricted this analysis to singleton variants. For each trait, we estimated the sampling-error variance from synonymous variants in genes not significantly associated with that trait, and assumed it to be the same for all variant categories. We then defined the detectable variance for each variant category as the observed variance in excess of this estimated shared sampling-error variance. Across 444 genes with at least 10 singleton missense variants both within and outside TED-annotated domains and scored by all methods, the detectable variance was on average 6.6% (95% CI [4.7, 8.6], across 2,000 gene bootstraps) of the observed variance within TED-annotated domains and 3.0% (95% CI [1.7, 4.5]) outside them. Thus, both regions contained detectable between-variant variation, with a larger spread of effects within structured regions. Across all missense variants, the detectable variance was on average 5.0% (95% CI [3.7, 6.3]) and the ranking of methods by captured variance was very similar to that observed within TED-annotated domains, although with lower values of captured variance (Supplementary Fig. 9). For each score, we used linear regression within each gene-trait pair and region to quantify the variance captured after correcting for sampling error (Fig. 4c,d). Unlike the rank-based Spearman correlations used elsewhere, this quantity reflects how well a linear relationship with the score explains variation in variant effect estimates, rather than only variant ordering. CPT-1, AlphaMissense, popEVE, AVI, ESM1v, and BayesDel explained roughly 20% of the detectable variance within TED domains, with overlapping intervals, while conservation scores explained at most 13.4% (95% CI [8.8, 18.8]), and the gnomAD allele-frequency explained none. The same methods, except for AVI, appeared to lead outside TED domains. Although these regions are statistically underpowered to establish firm rankings, our results indicate that the best-performing methods leave most of the detectable between-variant variation unexplained, even within structured protein domains. A part of this uncaptured variance may reflect the linear fit rather than ranking error, since a score that ranks variants perfectly need not relate linearly to the effect estimate.

Among indels, LOFTEE HC status explained 26.2% (95% CI [17.1, 37.5]) of the detectable variance, and indel length about 13.9% (95% CI [8.4, 20.0]), while the allele-frequency baselines explained at most 6.4% (95% CI [1.1, 12.0]; Fig. 4e). Predicted loss-of-function variants are usually treated as interchangeable and collapsed into a single burden per gene ^40,66^. However, LOFTEE HC variants themselves showed substantial detectable variance (7.8%; 95% CI [5.1, 10.6]), of which the additional variant scores examined explained only a small fraction (Fig. 4f). Under the assumption that loss-of-function variants affect the phenotypic mean but not its variance, these results suggest that predicted loss-of-function variants may differ appreciably in their phenotypic effects and that much of this heterogeneity is not captured by the variant scores examined here.

Altogether, UKBBGym is reproducible from publicly available summary statistics and allows assessment of how well variant scores capture phenotypic variation.

### Experimental assays do not consistently outperform computational scores in predicting missense effects in human carriers

Deep mutational scanning (DMS) assays measure variant effects in molecular or cellular systems, whereas population phenotypes capture the consequences of these variants in human carriers. We therefore asked how closely variant effects inferred from experimental readouts and carrier phenotypes agree, both for individual variants and in their evaluation of computational predictors.

We identified four genes with sufficient overlap between ultra-rare variants, functional measurements, and associated traits. For *LDLR*, cellular readout of LDL uptake ^71^ correlated with variant effect estimated on LDL cholesterol from human carriers whereas cell-surface abundance showed little correspondence, and twelve of seventeen computational scores correlated significantly more strongly than either experimental readout (Fig. 5a). For glucokinase (*GCK*), which regulates blood glucose by controlling glucose-stimulated insulin secretion, ten of seventeen computational scores correlated more strongly with variant effect estimated on HbA1c from human carriers, a marker of long-term blood glucose levels, than measured effects on *GCK* activity and abundance in humanized yeast-based assays ^72,73^ (Fig. 5b). For the lipid phosphatase tumour suppressor *PTEN* ^74^ and the adipocyte differentiation regulator *PPARγ* ^75^, experimental and computational scores showed broadly comparable correlations with variant effect estimates based on their associated traits (Fig. 5c,d). Together, these data indicate that high-resolution functional measurements do not necessarily predict human carrier phenotypes better than computational variant scores.

**Figure 5:**
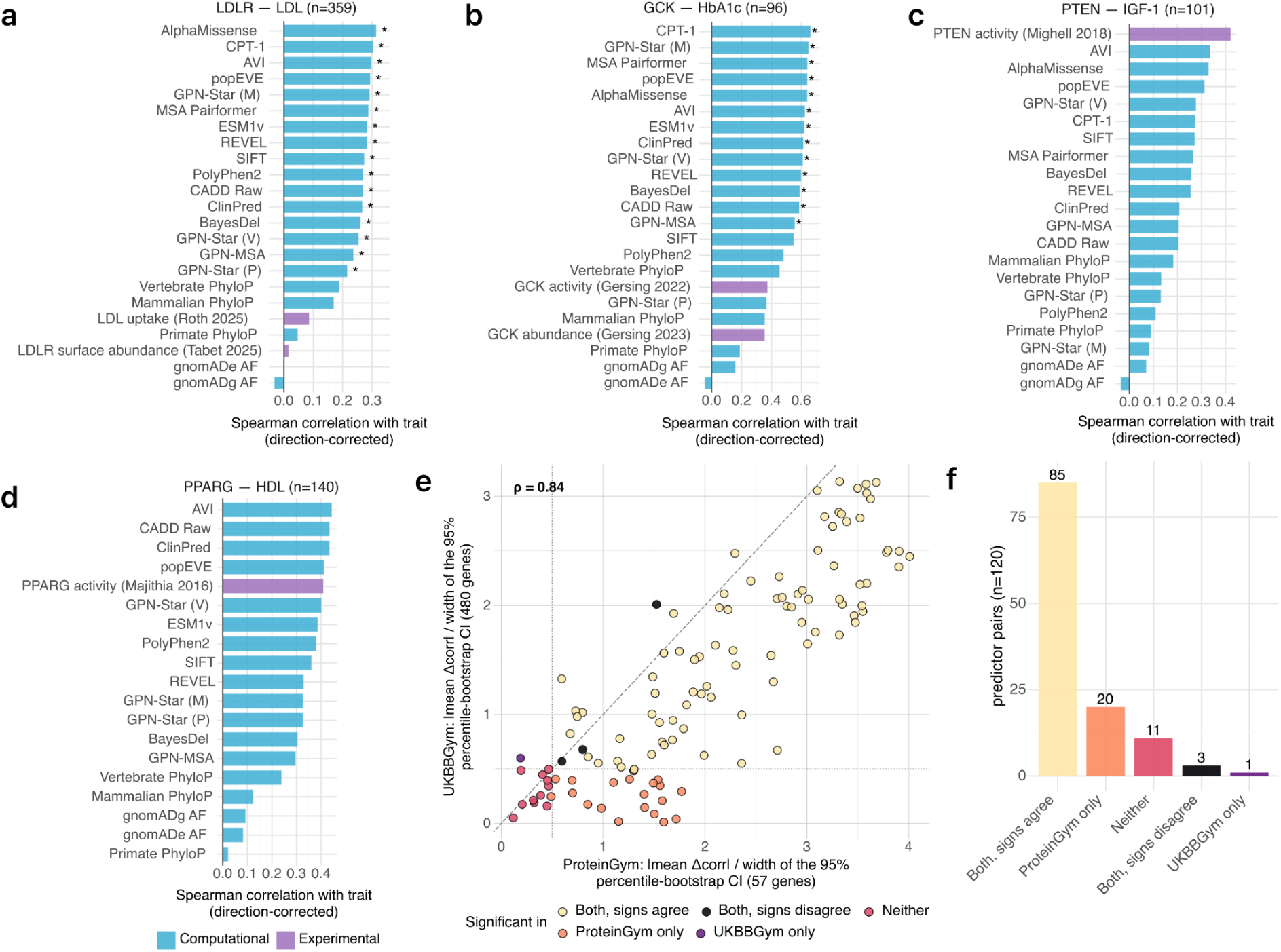
Experimental assays do not consistently outperform computational scores for missense effects in human carriers. **a**, Spearman correlation of computational variant scores and experimental functional scores with LDL cholesterol effect estimates for 359 *LDLR* missense variants. Experimental scores include *LDLR* cell-surface abundance and LDL uptake ^71^. * indicates computational scores whose correlations are significantly greater than that of all experimental scores (*P* < 0.05, based on 10,000 bootstraps). **b,** As in (a) for glycated hemoglobin (HbA1c) across 96 *GCK* missense variants. Experimental scores include *GCK* enzymatic activity ^72^ and *GCK* protein abundance ^73^. **c,** As in (a) for IGF1 across 101 *PTEN* missense variants. Experimental scores are from the *PTEN* lipid phosphatase activity assay ^74^. **d,** As in (a) for HDL cholesterol across 140 *PPARγ* missense variants. Experimental scores are from the *PPARγ* CD36 expression assay ^75^. **e,** Signal-to-noise ratio comparison in ProteinGym versus UKBBGym (implemented using Genebass). For each pair of variant scores, the signal-to-noise ratio is defined as the mean difference in correlation divided by the width of its 95% bootstrap confidence interval, across 10,000 bootstrap samples. Point color indicates whether each pair is significant and sign-concordant across the two benchmarks. **f,** Number of variant score pairs for categories defined in (e).

We next asked whether UKBBGym and ProteinGym ^31^, a compendium of DMS datasets, resolve differences between computational predictors similarly. While correlations were consistently higher in ProteinGym than in UKBBGym, consistent with noisier population-level phenotype measurements and denser missense variant measurements in DMS assays, the relative ranking of variant scores was broadly preserved (Supplementary Fig. 10). Accordingly, signal-to-noise ratios were higher in ProteinGym for most pairwise score comparisons, indicating greater power to resolve differences between scores (Fig. 5e). Nonetheless, 85 score pairs were significantly distinguished in both benchmarks and agreed in the direction of the difference, whereas 20 were significant only in ProteinGym, and 1 (between GPN-Star (M) and GPN-MSA) was significant only in UKBBGym (Fig. 5f). Three score pairs were significant in both benchmarks but disagreed in direction: on UKBBGym, AVI outperformed ESM1v and REVEL, and Mammalian PhyloP outperformed Primate PhyloP. Thus, despite its lower resolving power, UKBBGym recovers the vast majority of the same pairwise differences while using human carrier phenotypes.

### Population-phenotype performance tracks clinical classification without favoring scores trained on clinical labels

We next asked how variant effect estimates based on population phenotypes relate to established clinical variant classifications. Stratifying variants by ClinVar annotations, pathogenic and likely pathogenic variants showed the lowest average effect on protein abundance and the largest trait effects, whereas benign and likely benign variants showed mean estimated effects near zero for quantitative traits and protein abundance (Fig. 6a). This shows that UKBBGym captures phenotypic differences corresponding to clinical classification while retaining a quantitative measure of variant-associated phenotypes.

**Figure 6.**
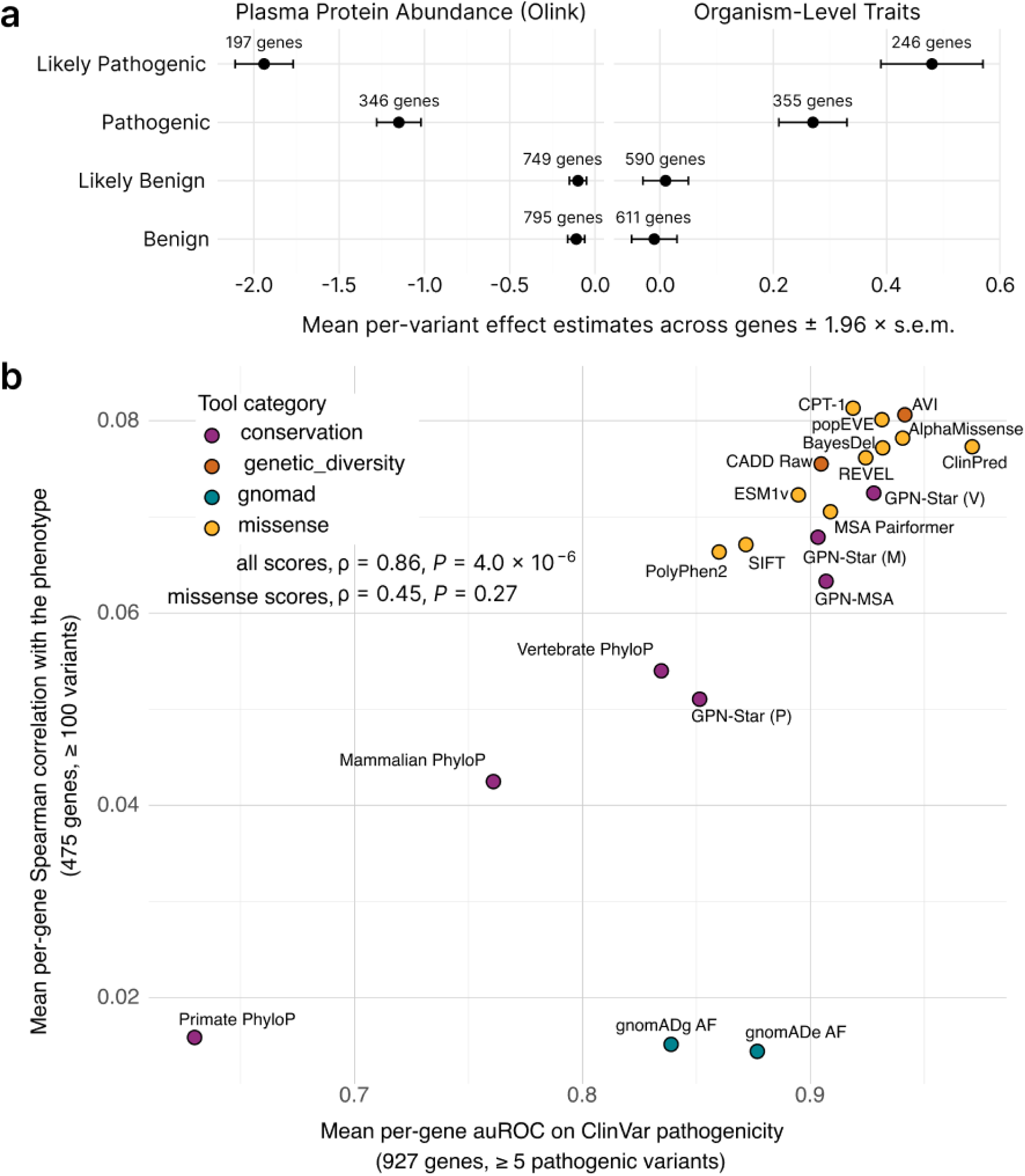
Population-phenotype performance tracks clinical classification without favoring scores trained on clinical labels. **a**, Mean across genes of estimated variant effects (left: protein abundance, right: traits) averaged within each gene, for missense variants stratified by ClinVar clinical significance classification. Error bars indicate 1.96 × s.e.m. **b,** Mean per-gene Spearman correlation between variant scores and variant effect estimates (y-axis; 475 gene-trait pairs for genes with ≥ 100 missense variants) against mean per-gene auROC for ClinVar clinical significance classification (x-axis; 927 genes with ≥ 5 ClinVar-annotated pathogenic and ≥ 5 ClinVar-annotated benign variants). Each point represents a variant scoring method and is colored by functional category. Spearman correlation (ρ) across all scores and across missense scores is indicated. *P* values obtained across 100,000 permutations.

Intriguingly, likely pathogenic variants were associated with significantly larger effects than pathogenic variants (Fig. 6a) even when restricting the analysis to the same genes (Supplementary Fig. 11a). Likely pathogenic variants were also rarer in the cohort than pathogenic variants (Supplementary Fig. 11b), suggesting that allele count may contribute to this ordering. One possible explanation is ascertainment in clinical classification: recurrent variants may accumulate sufficient evidence to be classified as pathogenic through independent observations of the same variant in unrelated affected individuals ^27^, whereas even rarer variants with comparable or stronger effects may remain classified as likely pathogenic. Consistent with an allele-count contribution, the difference between pathogenic and likely pathogenic variants was no longer significant when both groups were restricted to lower allele counts (Supplementary Fig. 11c) ^27^.

Next, we compared how methods rank missense variants for ClinVar classification versus phenotypic effects in UKBBGym. Since ClinVar has limited coverage of non-coding regions (only 752 pathogenic or likely pathogenic non-coding variants outside canonical splice sites), we focused this analysis on missense variants. Across methods, the ability to discriminate ClinVar pathogenicity positively correlated with their performance on UKBBGym (ρ = 0.86, *P* = 4.0 × 10^-6^; Fig. 6b), showing that population-phenotype performance broadly tracks clinical utility. Because allele frequency from gnomAD is often used as evidence for benign classification in ClinVar ^27^, gnomAD exome and genome frequencies distinguished ClinVar labels comparably to several sequence-based predictors. Restricting UKBBGym to ultra-rare variants leaves gnomAD allele frequencies with little dynamic range for ordering variants, resulting in poor performance on UKBBGym. Among missense predictors alone, however, the two benchmarks were not significantly correlated (ρ = 0.45, *P* = 0.27), showing that they resolve the leading predictors differently (Fig. 6b). ClinPred, which was trained to recover clinical labels, achieved the highest within-gene median auROC for ClinVar classification, but was less effective at ordering variants in UKBBGym than models not trained on clinical labels. ^54^, In contrast, CPT-1, trained using deep mutational scanning data, performed relatively better in UKBBGym than on ClinVar. This divergence is consistent with known circularity in benchmarks based on clinical labels, which can favor predictors trained on the same or related data ^54,76^.

Altogether, UKBBGym ranks variant scores in agreement with clinical classification while remaining independent of clinical labels, making it suited to assessing and developing predictors for clinical use.

## Discussion

In conclusion, UKBBGym establishes population-scale human phenotypes for evaluating coding and non-coding variant scores. UKBBGym recapitulates relative performance on pathogenicity prediction without suffering from legacy biases in clinical variant databases and potential mismatches between experimental assay readouts and quantitative phenotypes. Owing to its breadth in variant and phenotypic modality coverage, UKBBGym reveals stronger associations with protein abundance and quantitative traits for scores modeling effects on the protein product than for those modeling transcriptional regulation. Comparing captured to detectable variance indicates room for improvement for various variant categories, notably among indels and loss-of-function variants. Finally, UKBBGym can be reproduced using publicly available summary statistics, thereby lowering the barrier to evaluating the scores of coding variants against human phenotypes for researchers without access to individual-level data.

UKBBGym provides an alternative to experimental benchmarks for evaluating and disentangling clinical label bias, while retaining the advantage of being grounded in quantitative human phenotypes. We found that ProteinGym generally exhibited a higher signal-to-noise ratio than the Genebass summary-statistics implementation used in UKBBGym. However, for most comparisons, this difference did not substantially affect the conclusions. Importantly, a higher experimental signal does not necessarily imply better correspondence with quantitative phenotypes. For the four proteins for which a direct comparison was possible, we found that MAVE readouts and observed human phenotypes were not always well aligned, as illustrated by *LDLR* and related examples. This highlights a broader challenge in translating measurements from in vitro assays to effects in vivo. Similar limitations in the translatability of experimental models are well recognized across biomedical research and contribute to the failure of some therapeutic candidates when moving from preclinical studies into clinical development ^77^. As MAVEs become available for a larger number of trait-associated genes, systematic comparisons with population-scale human genetic phenotypes could therefore help identify which experimental assays most faithfully capture consequences in human carriers and guide the development of assays with improved physiological relevance.

An important purpose of benchmarking is to indicate where improvements are needed and worth pursuing. Relative to the variance detectable given the precision of estimated variant effects, the best missense-variant predictors accounted for up to 30% of that variance, indicating substantial room for improvement. This analysis relies on Pearson correlation and therefore rewards models whose scores are linearly related to the phenotype, without prioritizing variants with extreme effects. Hence, part of the missing variance may be captured by a non-linear, monotonic mapping of scores to traits, as recently proposed for burden testing ^78^. Moreover, independent analyses of detectable variance for other variant-scoring benchmarks would be valuable. For the multiplexed assay community, for instance, routine reporting of per-variant measurement variation would allow computation of an analogous ceiling ^79^.

Our benchmark has limitations. As a cohort of largely healthy volunteers, the UK Biobank is depleted of highly penetrant alleles, and the disease-causing variants it contains are observed only in heterozygous carriers, precluding the modeling of dominant and recessive effects. Our analysis is further restricted to protein-coding genes, excluding non-coding genes and other intergenic regulatory elements, and to SNVs: insertions and deletions are infeasible to score at this scale for most sequence-based models.

Evaluating predictors of variant deleteriousness will remain important for deciding which methods to use in rare disease diagnostics, cancer variant interpretation, and burden testing. Extending UKBBGym to further molecular modalities and cohorts is a natural next step, and unlike curated label sets, which grow only with expert annotation, population phenotype data accumulates passively with cohort size and sequencing depth. As biobanks grow, we expect benchmarks built on such phenotypes to play a larger role alongside clinical labels and experimental assays in evaluating variant effect predictions.

## Methods

### Data and cohort

#### Sample selection

We restricted the analysis to samples listed under UKB field 22006, which defines samples that self-identified as ’White British’ and have very similar genetic ancestry based on principal components analysis of the genotypes.

We performed kinship-based pruning using a greedy algorithm adapted from ukbtools ^80^ and reimplemented in Python. Among individuals with available phenotypic data, we iteratively removed a random individual from those with the most relatives at third degree or closer (i.e., kinship > 0.0884) until no pairs exceeded the kinship threshold, resulting in 379,645 individuals.

#### WGS Genotype processing

Single-nucleotide variants (SNVs) and short indels from UK Biobank whole-genome sequencing, jointly called using DRAGEN 3.7.8, were retrieved from the DRAGEN pVCF files provided by UKB (Field 24053) ^38^. Following refs. ^80,81^, using bcftools ^82^ (v1.23), we considered genotype calls missing if the sum of local allele depths (sum(LAD)) was <8 or the genotype quality (GQ) was <10 per sample genotype. We subsequently dropped any variant with a missingness >10%. Finally, using bcftools (v1.23), multi-allelic sites were split, and indels were normalized and left-aligned based on a GRCh38 reference assembly available at https://ftp.ebi.ac.uk/pub/databases/gencode/Gencode_human/release_40/GRCh38.primary_assembly.genome.fa.gz. This genotype QC was applied across 9,843 pVCFs covering the genes used in this study. Non-reference genotypes were extracted into a sparse format of sample ID, variant ID, and genotype encoded as 1 for heterozygous or 2 for homozygous alternate, and written to parquet files restricted to variants with allele count below 20. Unique variants were additionally stored in a companion metadata parquet file for subsequent variant annotation. Indels longer than 50 bp were excluded from all variant-level analyses. Throughout this study, we considered variants with an allele count ≤ 20 to be ultra-rare, unless otherwise stated.

#### Variant annotation

Unique variants were annotated using the Ensembl Variant Effect Predictor ^83^ (VEP, version v113) against the GRCh38 reference assembly and Gencode release 40 gene models ^84^, with the LOFTEE plugin ^41^ for loss-of-function variant classification. VEP was run with a single canonical transcript per variant selected using a priority-ordered pick strategy: transcript biotype, MANE Select ^85^, canonical status, APPRIS ^86^, transcript support level, CCDS, consequence rank, transcript length, and finally Ensembl, then RefSeq source. All variants within ±5 kb of the gene body of a protein-coding gene were annotated to their respective genes. Only protein-coding genes were retained. VEP provided variant consequences on the MANE Select transcript, gene, and protein coordinates, distance to the transcription start site, SIFT and PolyPhen-2 scores, gnomAD allele frequencies from both genomes and exomes, and LOFTEE high and low confidence annotations ^41^. We additionally derived variant length and indel status (indel status was decomposed into three binary indicators: is_indel, is_deletion, and is_insertion).

Additional annotations were computed separately and merged with the VEP output by variant identifier. GPN-MSA ^11^, retrieved by remote tabix ^87^ from precomputed genome-wide scores. CADD v1.7 raw scores ^22,53^ were retrieved the same way from the GRCh38 whole_genome_SNVs table for SNVs and the gnomad.genomes.r4.0 table for gnomAD indels. PhyloP ^6^ scores from the vertebrate 100-way, mammalian 17-way, and primate 30-way alignments, read from the corresponding UCSC bigWig tracks. Precomputed AlphaGenome Variant Impact (AVI) ^23^ and GPN-Star Vertebrate, Mammalian, and Primate ^12^ scores were obtained for all SNVs.

For splicing, we used the maximum AbSplice ^18^ across tissues. We also used SpliceAI ^16^ and Pangolin ^17^ delta scores, extracted from the precomputed AbSplice score files, respectively. For expression and regulation, we used the minimum AbExp ^48^ score across tissues and the PromoterAI ^14^ score and its absolute value. We also used Flashzoi ^13,47^ (replicate 0) predictions for SNVs in predefined promoters and enhancers. For each RNA-seq track, the predicted variant effect was defined as the difference in log1p-transformed exon-summed coverage between the alternative and reference alleles. We report the minimum predicted effect across either all RNA-seq tracks or the subset corresponding to GTEx tissues (track IDs 7522–7610). Precomputed missense variant scores comprised AlphaMissense pathogenicity ^20^, REVEL ^46^, ClinPred ^50^, BayesDel ^52^, the CPT-1 ^44^, ESM1v ^45^, and popEVE ^62^. We computed MSA Pairformer ^9^ log-likelihoods computed from AlphaFold DB MSAs. Missing scores were set to each method’s neutral value, as listed in Supplementary Table 3.

Candidate cis-regulatory elements (cCREs) were obtained from the ENCODE SCREEN registry ^61^, which annotates each element by regulatory class. Variants were assigned to overlapping cCREs by genomic interval overlap and represented by per-class binary indicators for promoter-like (PLS), proximal and distal enhancer-like (pELS and dELS), chromatin-accessible (CA), CA-CTCF, CA-TF, CA-H3K4me3, and TF-only elements, together with a residual ‘not annotated in ENCODE’ category.

Protein-domain membership was annotated for the Ensembl-canonical protein of each protein-coding gene using three sources: MobiDB ^58^ for intrinsically disordered regions (selecting regions with source “curated”, type “disorder”, and subtype “priority”) and linear interacting peptides (selecting regions with type “lip” and subtype “priority”), TED ^57^ predicted structural-domain boundaries, and low-complexity domains obtained from LCD-Composer ^59^. The protein positions of each variant were mapped to these domain intervals to produce binary indicators (structural domain, disordered region, linear interacting peptide, low-complexity domain). Per-residue structural confidence was also annotated using the AlphaFold2-predicted local distance difference test (pLDDT) scores (AlphaFold human proteome, v6) ^70^, with residues thresholded at pLDDT > 70 classified as structured and the rest as disordered.

Protein interfaces were annotated using two sources. High-confidence protein-protein interface residues were obtained from PIONEER ^68^. A second, independent interface annotation and per-residue inter-chain contact count (salt bridges, hydrogen bonds, and non-bonded contacts) were obtained from the Genomics2Proteins resource ^88^, from which binary “has-contact” indicators were derived.

#### Genomic region annotation

Region-type variant classes (enhancer, promoter, UTR, coding, intron) were each defined independently and were therefore not mutually exclusive. Hence, a variant can, for example, be both an intronic variant and an enhancer variant if it falls within an intron that also overlaps an ENCODE enhancer-like cCRE. Promoters were defined as the region within 500 bp of a transcription start site (TSS ± 500 bp) using Gencode release 39 to match the definition of PromoterAI. Enhancer variants were defined as those overlapping a proximal or distal enhancer-like signature cCRE (pELS or dELS) as defined above. Coding sequences in any annotated transcript and variants likely to affect splicing (Pangolin score ≥ 0.2) were removed from enhancer and promoter variants to better isolate gene-regulatory effects of these variants. For intronic variants specifically, we excluded coding sequences in any annotated transcript and core splice-site variants (i.e., those annotated as splice donor or acceptor sites by VEP).

To compare variant coverage across benchmarks, we tabulated variant counts the functional region assigned from VEP consequences and ENCODE cCRE annotations (Coding, UTR, Promoter, Enhancer, Introns, Intergenic) for i) UKBBGym, considering the ultra-rare variants in genes required by the quantitative traits and protein abundance benchmarks (see Subsection Benchmark construction below), ii) ClinVar ^24^ excluding variants of uncertain significance, iii) ProteinGym considering all human deep mutational scanning substitution and indel variants ^31^, and iv) the Mendelian- and complex-trait matched-negative test sets from TraitGym ^25^, downloaded from Hugging Face. For TraitGym, Promoter and Enhancer were assigned using the provided ENCODE cCRE annotations (PLS/PLS_flank as Promoter; pELS/dELS and their flanks as Enhancer).

#### Covariates processing

We retrieved genetic sex (Field 22001), age at recruitment (Field 21003), and the first 40 genetic principal components (Field 22009), and derived age², age × sex, and age² × sex as additional covariates.

#### Quantitative traits

##### Quantitative traits processing

For measurements taken multiple times during a single visit (e.g., blood pressure readings), we used the average of all readings to produce a single per-visit measurement. Each phenotype was then transformed to approximate normality using a rank-based inverse normal transformation (RINT) with Blom’s correction (c = 3/8):

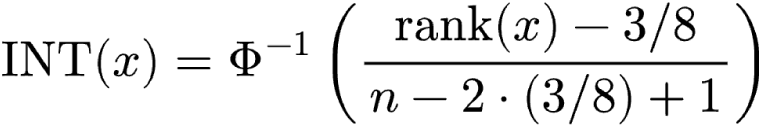

where *n* is the number of non-missing observations for a given phenotype and Φ^-1^ denotes the probit (inverse standard normal) function.

To adjust for common-variant contribution, we used REGENIE ^42^ (v4.12) step 1 with a block size of 1,000 variants and the print-prs flag to obtain a whole-genome polygenic prediction. Step 1 was run on directly genotyped autosomal array calls from UK Biobank data field 22418, merged and filtered with PLINK2 ^89^ to a minor allele frequency above 1%, a minor allele count above 100, a genotyping rate above 0.90, a Hardy-Weinberg equilibrium test *P* value above 10⁻¹⁵, and sample missingness below 0.1, then pruned at a pairwise linkage-disequilibrium r² threshold of 0.9 with a window size of 1,000 markers and a step size of 100 markers. We then regressed the RINT-transformed phenotypes on this polygenic score and the covariates, using the residuals as the final phenotype values for downstream analyses.

##### Gene-trait pairs

For each protein-coding gene, we constructed a per-individual burden score as the count of ultra-rare LOFTEE HC (high-confidence loss-of-function) variants each individual carried in that gene. We used REGENIE step 1 to generate leave-one-chromosome-out (LOCO) predictions from imputed array genotypes. Burden scores were then passed as BGEN files to REGENIE step 2, testing for association with RINT-transformed phenotypes (--apply-rint), conditional on the LOCO predictions from step 1 (block size of 1,000 variants for both steps). Association testing was performed across 112 quantitative phenotypes with less than 20% missingness. Gene-trait pairs were declared significant at a false-discovery rate ≤ 0.05 (Benjamini-Hochberg ^90^, correcting across all gene-trait pairs, Supplementary Table 1).

For each gene, we retained the trait with the largest absolute Spearman correlation (LOFTEE HC status vs. estimated variant effect) among its significant associations, yielding the 670 gene-trait pairs considered throughout this benchmark across 96 traits.

##### Direction of association

To orient the direction of each gene-trait pair consistently, we computed the Spearman correlation between the LOFTEE HC status of each variant and its estimated effect across all ultra-rare variants in the gene. This correlation captures whether predicted loss-of-function variants in a given gene tend to increase or decrease the phenotype. Gene-trait pairs for which the correlation could not be computed were excluded. Trait values were then oriented accordingly, so that variants with higher predicted impact were expected to be consistently aligned with higher trait values across all associations.

#### Olink plasma protein abundance

##### Olink plasma protein abundance processing

Olink NPX measurements from the UK Biobank initial assessment cohort (Field 30902 ^39^) were extracted, yielding ∼39,000 participants across ∼2,850 proteins when restricting to unrelated individuals of European ancestry. Quality control procedures, including outlier removal and a log_2_ transformation, had been applied prior to data release. To account for technical confounders, polygenic effects, and trans-regulatory effects on protein abundance, we evaluated several adjustment methods.

Our primary approach consisted of two steps. First, we estimated a per-protein polygenic prediction from the pruned array genotypes using REGENIE step 1 with the print-prs flag, with age, sex, diet, 40 genetic principal components, and 90 proteomic principal components as covariates. The polygenic score-corrected protein levels were then used as input for PROTRIDER ^43^ using the default settings, which aims at capturing trans-regulatory and other sources of shared variation. Model residuals were fit to a protein-specific Student’s t-distribution with per-protein location, scale, and degrees of freedom parameters. The residuals were then quantile-mapped to the standard normal distribution via the fitted t-CDF. The resulting z-scores constitute the adjusted protein levels.

As an alternative, we first corrected protein levels for polygenic effects, then regressed out known confounders of protein abundance. Specifically, the polygenic-corrected values across individuals were RINT-transformed, regressed on the same covariates listed above, and then RINT-transformed again, as in ref. ^14^. As a simpler baseline, we also computed per-protein z-scores across individuals.

##### Gene-plasma protein abundance pairs

To independently benchmark the protein adjustment methods, we selected ultra-rare variants likely disrupting protein abundance. These were defined as variants located within 5,000 bp of a gene and predicted to be loss-of-function (pLoF) by LOFTEE HC or likely pathogenic missense variants (AlphaMissense > 0.56 ^20^). For each method, we ranked the adjusted protein levels across all samples and, for each rank, estimated the proportion of samples with a variant likely disrupting protein abundance at or below that threshold. Since PROTRIDER consistently performed favorably in this benchmark (Supplementary Fig. 1), we adopted this adjustment for all subsequent analyses.

To ensure that plasma protein abundance provided an informative readout of loss of gene function, we retained only gene-protein pairs for which LOFTEE HC variants were associated with reduced abundance, following a similar strategy used in previous UK Biobank proteomic benchmarks ^36^. To this end, we tested the association between adjusted protein levels and the presence of a pLoF variant using a Wilcoxon rank-sum test, because the number of carriers per gene was often small. OLS regression was fit separately to estimate per-gene effects. We retained only genes with a significant Wilcoxon rank-sum test (false discovery rate < 0.05, Benjamini-Hochberg) and a negative OLS effect estimate, as pLoF variants are expected to reduce protein abundance, yielding 1,086 genes, of which 1,076 had at least 100 variants (Supplementary Table 2).

### Benchmark metrics

#### Spearman correlations

To assess the ability of variant scores to rank variants by phenotypic value, we computed the Spearman rank correlation between variant scores and estimated variant effects for each gene-phenotype pair and each tool. For some analyses, correlations were restricted to variants of a category of interest, e.g., missense variants. Pairwise comparisons between all annotation tools were performed using the two-sided Wilcoxon signed-rank test on Spearman correlations from intersecting gene-phenotype pairs between any two tools.

To test whether measured abundance could itself serve as a variant score, each variant’s carrier-mean adjusted Olink protein abundance was treated as a score and correlated with the estimated variant effects using the same Spearman procedure, across the 19 genes with matched Olink data and at least 100 missense variants.

#### Mean variant effect of top N variants

For each gene-phenotype pair (quantitative traits or protein abundance), variants were ranked in descending order of deleteriousness according to each score. The cumulative mean variant effect at rank *k* was defined as the average of the variant effects of the top-*k* variants in the gene. The cumulative mean variant effect as a function of rank was aggregated across all gene-phenotype pairs, yielding the mean ± s.e.m. for gene-phenotype pairs.

To handle variants with tied scores, we assigned each tied variant the mean estimated variant effect across all variants in the tied group. The tied variants were then assigned consecutive ordinal ranks, such that a variant effect was defined at every rank (k).

Pairwise comparisons between variant scoring methods were performed using the two-sided Wilcoxon signed-rank test on the cumulative mean phenotype at rank 5, restricted to genes present in both tools with at least ten matched pairs.

#### Mean variant effect by variant category

To compare phenotypic values across functional or clinical variant categories, variants were assigned to binary categories based on their functional annotations (e.g., ClinVar pathogenic or likely pathogenic, ClinVar benign or likely benign, structural domain membership, or indel length). Indel length bins were defined separately for insertions and deletions, each split into three mutually exclusive bins by length: 1 bp, 2-5 bp, and >5 bp. For each gene-phenotype pair, the mean estimated variant effect was computed across all variants in a given category, and then averaged across pairs.

## Benchmark using summary statistics

UKBBGym estimates effects of ultra-rare variants as the average zero-centered phenotype across its carriers. Published single-variant analyses instead report a regression coefficient (effect estimate) ^40,66^.

The average phenotype across carriers *θ*^ is a rescaling of the single-variant effect estimate *β*^ reported by standard association models, that is, the coefficient on *G* in the fit,

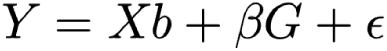

As detailed in Supplementary Note 1, by the Frisch-Waugh-Lovell theorem ^91,92^, the two estimators share the same numerator and differ only in their denominators. For *p* << 1, we approximately have:

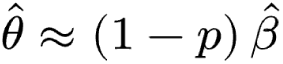

Where *p* is the carrier frequency of the variant. For UKBBGym variants, *p* is less than 5.1 × 10^-5^ (allele counts less than or equal to 20 and cohort sizes greater than 390,000), resulting in *θ*^ and *β*^ being almost identical for variants considered in UKBBGym.

### Genebass

To reproduce analyses for coding variants, encompassing predicted loss-of-function, missense, and synonymous variants, we used published single-variant statistics from Genebass, covering 394,841 UK Biobank exomes ^40^. Genebass single-variant summary statistics were downloaded from a publicly available Google Cloud Storage bucket: gs://ukbb-exome-public/500k/results/variant_results.mt.

Specifically, instead of using per-variant carrier means as the estimated variant effects (θ̂), we leveraged per-variant effect estimates from Genebass (using the field *BETA*), and recomputed the UKBBGym metric without further modification. The same gene-trait pairs as in the full UK Biobank analysis were used for the benchmark.

### All by All

GWAS and RVAS results from 392,030 individuals in the All of Us Research Program ^93^ (v8 release) were obtained from the All by All study ^66^. We used the cross-ancestry meta-analysis across two phenotype categories: physical measurements and laboratory measurements. A gene-trait pair was declared significant at an exome-wide threshold of *P* ≤ 6.7 × 10⁻⁷, using the meta-analysis field *META_Pvalue_Burden*. This resulted in 260 significant gene-trait pairs using a maximum minor allele frequency of 10^-3^. Only 76 genes met the minimum threshold of 100 missense variants required for the Spearman correlation analysis.

For every significant gene-trait pair from the All by All meta-analysis, single-variant statistics were extracted from the corresponding meta-analysis variant-level exome matrix tables, restricted to high-quality variants (hq_variant flag).

Variant-level metadata, i.e., the gene-specific most deleterious consequence on the canonical transcript, LOFTEE flags, PolyPhen and SIFT predictions, protein position, and per-ancestry allele frequencies, were joined from the All by All variant-QC-annotated table. Association direction was corrected using the field *META_Stats_burden*.

## Captured and detectable variance

Genebass single-variant effect estimates for singleton variants (allele count = 1) reflect both genuine between-variant differences in phenotypic effect and sampling error, i.e., phenotypic variability in single-variant carriers that is unrelated to the variant of interest.

As detailed in Supplementary note 2, we estimated the sampling-error variance 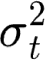 of each trait *t* from singleton synonymous variants in genes not significantly associated with that trait, which are expected to carry little systematic phenotypic effect and therefore isolate sampling error at the same allele-count and effect-size scales as the analysis variants.

To estimate the variance captured by a given score, we regressed the variant effect estimates on the variant score by ordinary least squares within each gene-trait pair and variant category. This compares scores on their linear association with the effect estimates rather than on their scale or sign. From the explained sum of squares of the regression, we subtracted its expected value when the score is unrelated to the true effects 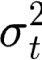 . Summing these over all gene-trait pairs and dividing by the summed degrees of freedom (the number of variants in each pair minus one) gives the captured variance of the score.

A perfect score, one that is an exact linear function of the true effects within each gene-trait pair and category, would leave only the sampling error unexplained, and the variance it captures is the detectable variance. We estimated this detectable variance by subtracting from the total sum of squares its expected sampling-error contribution, 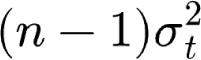, where n is the number of variants in each pair, summing over gene-trait pairs, and dividing by the summed degrees of freedom.

We report the variances with confidence intervals from resampling gene-trait pairs with replacement over 2,000 replicates and identical resampled indices across categories in paired comparisons.

### Analysis categories

For the within-versus-outside TED-annotated domains analysis, gene-trait pairs were retained only if they contained at least 10 variants within TED domains and at least 10 outside them, with finite scores from all methods compared. This yielded 444 gene-trait pairs and 70,356 variants.

For the missense analysis, we restricted to variants VEP annotated missense variants for all other methods compared. Gene-trait pairs with at least 10 qualifying variants were retained, yielding 555 pairs and 15,139 variants.

For the indel analysis, variants were required to be indels with a resolved indel-type annotation and finite values for all methods compared (Variant Length, LOFTEE HC, gnomAD exome/genome AF). Gene-trait units with at least 10 qualifying variants were retained, yielding 433 pairs and 10,880 variants.

For the pLoF analysis, we restricted to variants annotated as high-confidence loss-of-function (LOFTEE HC) for all other methods compared (Variant Length, CADD Raw, GPN-MSA, vertebrate/mammalian/primate PhyloP, gnomAD exome/genome AF). Gene-trait pairs with at least 10 qualifying variants were retained, yielding 481 pairs and 15,139 variants.

## Other variant score benchmarks

### Experimental assays comparison

#### Per-gene Spearman

We compared computational variant scores against experimental functional measurements, using their Spearman correlation with the Genebass effect estimate. Experimental data comprised ProteinGym (data downloaded from https://proteingym.org) deep mutational scanning substitution scores for human proteins ^31^ and gene-specific assays curated from the literature: *LDLR* cell-surface abundance and LDL uptake ^71^, *GCK* enzymatic activity ^72^, and protein abundance ^73^ (obtained via ProteinGym ^31^), *PTEN* lipid phosphatase activity ^74^ (obtained from ref ^76^), and *PPARγ* CD36 transactivation activity ^75^ (obtained via ProteinGym ^31^).

Target genes were mapped to Ensembl gene identifiers using each source’s identifier (UniProt accession or gene symbol), against the UniProt human ID-mapping file and the Ensembl-canonical protein-coding gene set. ProteinGym assay names, which use non-standard UniProt entry-name tokens, were first translated using a UniProt-derived lookup table (shared in the code repository for convenience and reproducibility; see Code availability).

Substitutions were matched to UK Biobank variants on gene, reference amino acid, protein position, and alternate amino acid. Where an assay provided multiple readouts for the same variant, each readout was retained separately.

We restricted this analysis to genes with an experimental assay overlapping at least 50 Genebass variants (*GCK* had the smallest overlap among included genes, with 96 variants; the next gene, *NPC1*, had 44). We computed the Spearman correlation of each computational or experimental score with the estimated variant effects.

To compare computational scores against experimental scores within each gene, we bootstrapped variants with replacement 10,000 times and recomputed each score’s Spearman correlation with the estimated variant effects. For each ordered pair of scores (*i*,*j*), we calculated the one-sided bootstrap *P*-value as the number of bootstrap iterations in which *ρ_i_* ≤ ρ_j_, plus one, divided by 10,001 ^94^. Each score was considered significantly better than all scores in the other category (computational tool vs. experimental assay) if its correlation significantly exceeded that of every score in the opposite category (*P* < 0.05). This variant-level bootstrap is distinct from the gene-level resampling used for the signal-to-noise analysis below.

For compatibility, we restricted the ProteinGym results to SNVs even though some methods report scores at the amino acid level.

#### Signal-to-noise ratio

To assess the discriminative power of each benchmark, that is, its ability to resolve meaningful differences between predictors, we computed, for each predictor, the Spearman correlation between the variant score and the phenotype across the variants of each gene. This was done for the 480 genes of UKBBGym and for the 57 human proteins assayed in ProteinGym. For each benchmark, we then bootstrapped the underlying set of genes 10,000 times, recomputing correlation on each resampled set for every predictor. For each pair of scores, at each bootstrap iteration, we recorded the difference in their correlations. For each pair, we defined the signal-to-noise ratio as the mean differential correlation divided by the width of the 95% equi-tailed interval across the 10,000 bootstrap samples.

#### ClinVar comparison

##### Per-gene auROC

ClinVar variant pathogenicity labels were downloaded on 2026-03-15 (VCF, GRCh38) from https://ftp.ncbi.nlm.nih.gov/pub/clinvar/vcf_GRCh38/ ^24^, and annotated with the same variant-annotation pipeline used throughout this study. We defined the positive class as the Pathogenic and Likely Pathogenic, and the negative class as the Benign and Likely Benign. Variants matching neither label were excluded, as were the small number of variants matching multiple labels. Genes were retained for this comparison only if they had at least 5 positive and 5 negative variants. For each retained gene and each annotation tool, scores were ranked within the gene using average ranks (tied scores receive the mean of the ranks they span), and the area under the receiver operating characteristic curve (auROC) was computed from the rank-sum identity, i.e.:

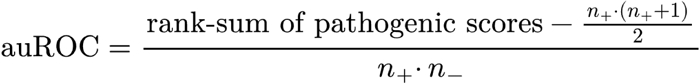

where *n*_+_ and *n*_-_ refer to the number of positive and negative variants, respectively. Per-gene auROC values were averaged across genes to obtain one summary auROC per tool. Genes were additionally required to have a score from every tool in the set common to both the ClinVar auROC and UKBBGym gene-wise Spearman-correlation comparisons, so that all tools are compared on the same gene set in both analyses.

#### Allele-count stratification

To assess whether ClinVar Likely Pathogenic variants having larger variant effect estimates than Pathogenic variants was related to differences in allele count, we compared Genebass variant effect estimate between Likely Pathogenic and Pathogenic variants across all gene-phenotype pairs containing LOFTEE HC, ClinVar Likely Pathogenic, and ClinVar Pathogenic variants, yielding 203 matched pairs. This comparison was repeated at progressively stricter allele-count thresholds (≤20, 5, 3, 2). Significance was assessed using a paired two-sided t-test.

## Data availability

The data to run the Genebass version of the benchmark is available on Zenodo (https://zenodo.org/records/22826446) ^95^ and on HuggingFace (https://huggingface.co/datasets/gagneurlab/ukbbgym).

## Code availability

Code used to generate the UKBBGym benchmark datasets and recreate the results is available at https://github.com/gagneurlab/ukbbgym.

Code used for the preprocessing of samples, genotypes, quantitative traits, and Olink plasma proteomics is available at https://github.com/gagneurlab/ukbb-utils-gagneur.

DNA nexus applets for processing the data and running the benchmark will be made available to approved UK Biobank researchers upon re-opening of the UK Biobank.

## Author information

### Contributions

S.L and J.G. conceived the study. S.L and E.H. implemented the framework.

E.H. led UKBB data processing together with A.S., S.L., and G.T. G.T. led the Olink data processing. P.T.d.S made additional conceptual contributions to the method. J.H. ran Flashzoi. S.L., E.H. and J.G. wrote the manuscript with input from all the authors. H.F. conceived the summary statistics implementation. J.G. supervised the project.

## Acknowledgements

The authors would like to thank Nils Wagner, Oliver Stegle, Brian Clarke, Iva Pritišanac, Agnes Toth-Petroczy, Thomas Opsomer, and Konrad Karczewski for helpful discussions. This research has been conducted using the UK Biobank Resource under Application Number 81358. This study used data from the All of Us Research Program’s Controlled Tier Dataset (CDR v8), available to authorized users on the Researcher Workbench.

This work was supported by the Initiative and Networking Fund of the Helmholtz Association in the framework of the Helmholtz AI project call (project DeepVar, ZT-I-PF-5-156), the Deutsche Forschungsgemeinschaft (DFG, German Research Foundation) via the project TRIUMPH - Tracking and Reconstructing Interactions to Understand the Missing Parts of Heritability (535971044 to A.S.), and the European Research Council (101118521, EPIC to J.H. and J.G.). S.L. and E.H. were supported by the Helmholtz Association under the joint research school Munich School for Data Science - MUDS. S.L. was supported by the EMBO Scientific Exchange Grant (#12502). P.T.d.S. was supported by the Munich Center for Machine Learning. G.S. was supported by the project ERDERA from the European Union’s Horizon Europe research and innovation program under grant agreement N°101156595. Views and opinions expressed are those of the author(s) only and do not necessarily reflect those of the European Union or any other granting authority, who cannot be held responsible for them. The IT infrastructure at the Chair of CMM (TUM) was co-funded by the Deutsche Forschungsgemeinschaft (DFG, German Research Foundation) under Project IDs 461264291 and 553375143.

