## Supplementary Figures for "Phenotypes of ultra-rare variant carriers benchmark variant effect scores"

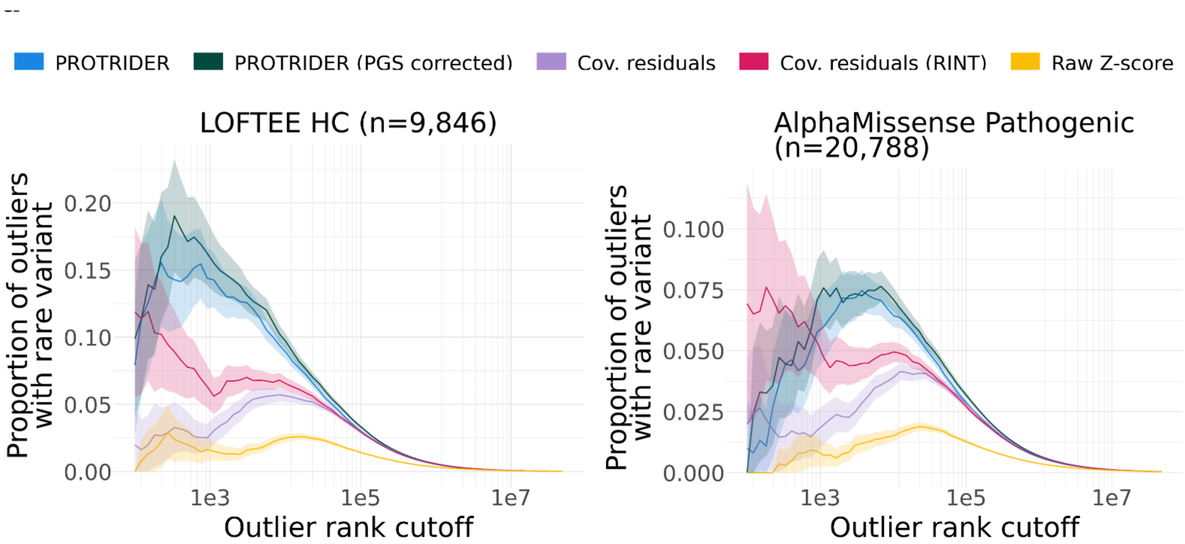


**Supplementary Figure 1: Benchmarking of protein abundance adjustment methods.** Proportion of samples (y-axis) at or below a given outlier rank cutoff (x-axis) that carry an ultra-rare predicted loss-of-function variant, annotated by LOFTEE HC (left) or AlphaMissense (right). Adjustment methods compared (see Methods): PROTRIDER, PROTRIDER (PGS corrected), Covariate regression residuals, Covariate regression residuals (RINT), and Raw Z-score. Ribbons mark 95% confidence intervals.

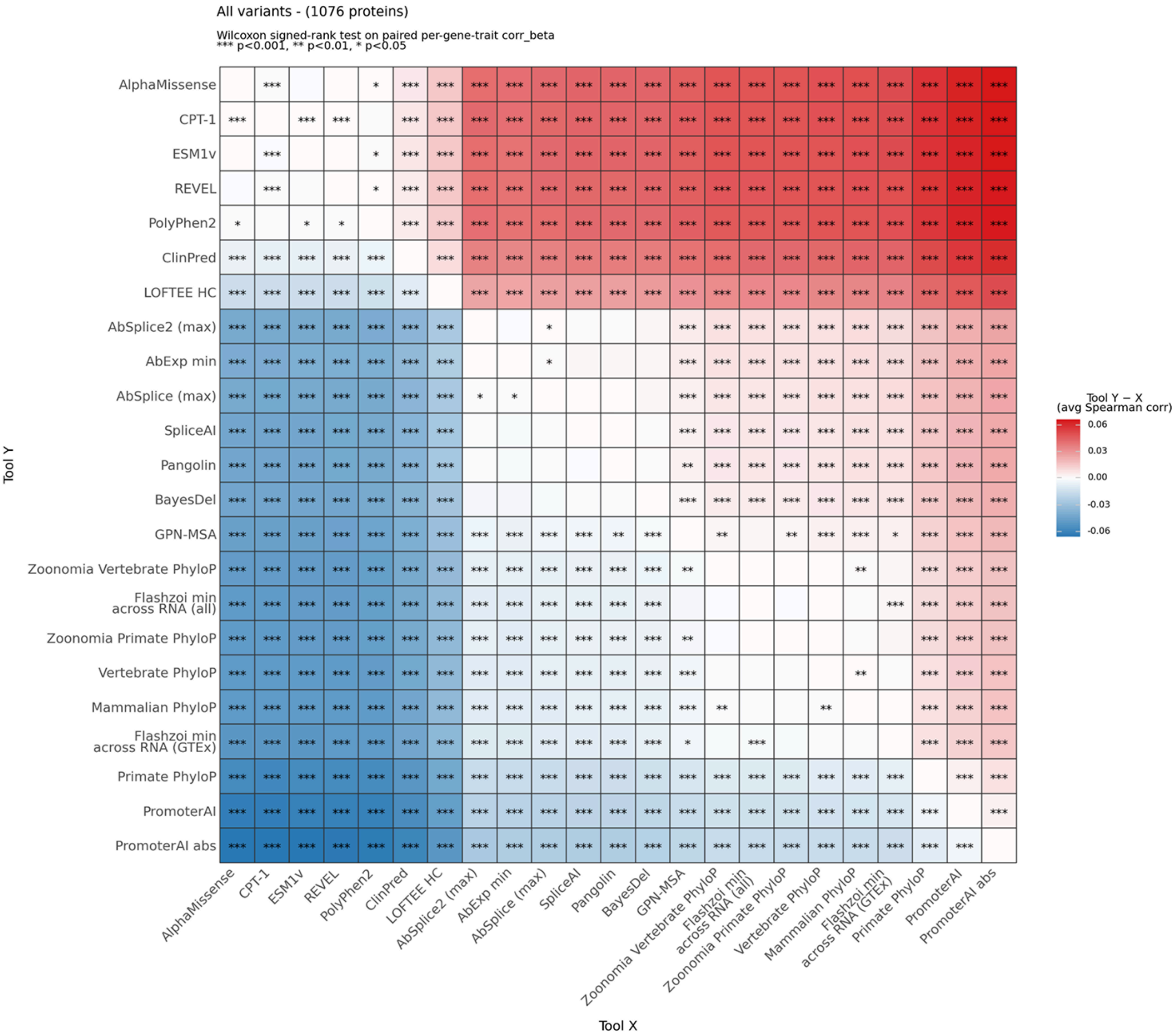


**Supplementary Figure 2: Pairwise comparison of variant-scoring methods on plasma protein abundance.**

Pairwise comparison of Spearman correlations between all variant-scoring methods and variant effect estimates on plasma protein abundance across 1,076 proteins, for all variants. Each cell compares the row method (Tool Y) against the column method (Tool X): color shows the mean paired difference in Spearman correlation (Tool Y − Tool X). Red indicates the row method correlates more strongly with estimates of effect on abundance than the column method, and blue indicates the reverse. Significance of the paired correlations assessed by the Wilcoxon signed-rank test across proteins (*** *P* < 0.001, ** *P* < 0.01, * *P* < 0.05).

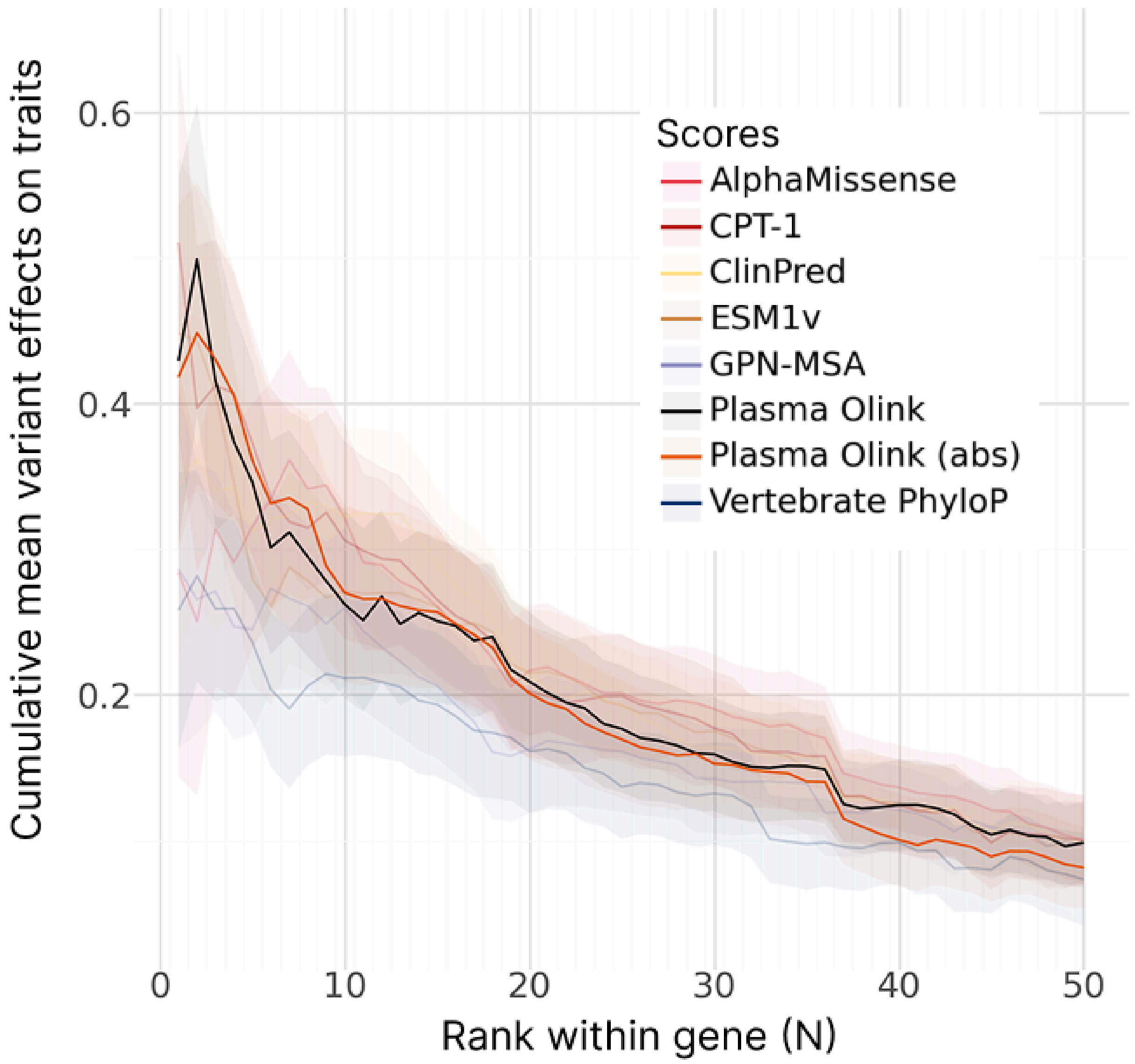


**Supplementary Figure 3: Variants associated with extreme plasma protein abundance are also associated with extreme trait values.**

Cumulative mean variant effect on traits (y-axis) across the top-ranked variants (x-axis) according to various scores for quantitative traits across 57 genes with Olink plasma protein abundance measurements. Gene-trait associations are direction-corrected such that loss-of-function variants correspond to an increase in phenotype value. Ribbons mark 1 × s.e.m. across genes.

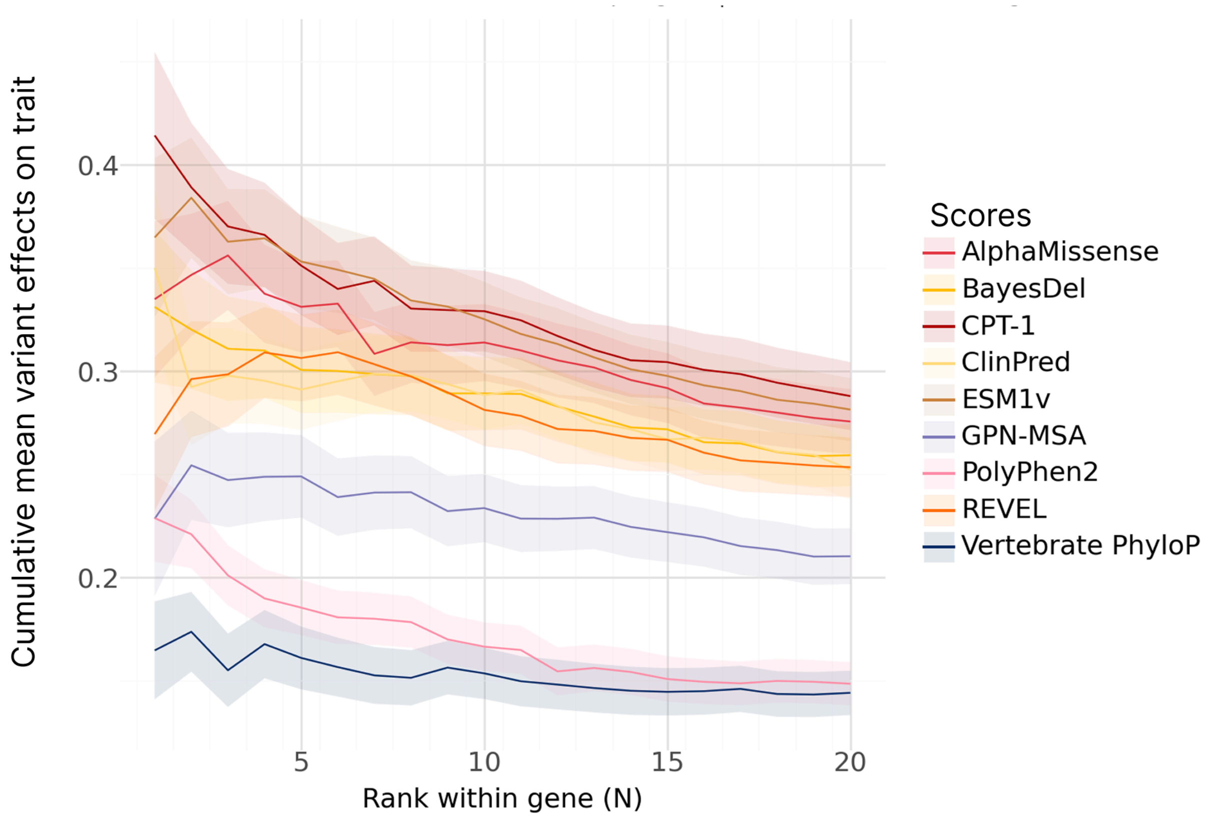


**Supplementary Figure 4: Cumulative mean variant effect estimates across the top-ranked variants by methods**

Cumulative mean variant effect estimates for quantitative traits (y-axis) across the top-ranked variants (x-axis) according to various scores for missense variants. Gene-trait associations are direction-corrected such that loss-of-function variants correspond to an increase in phenotype value. Ribbons mark 1 × s.e.m. across genes.

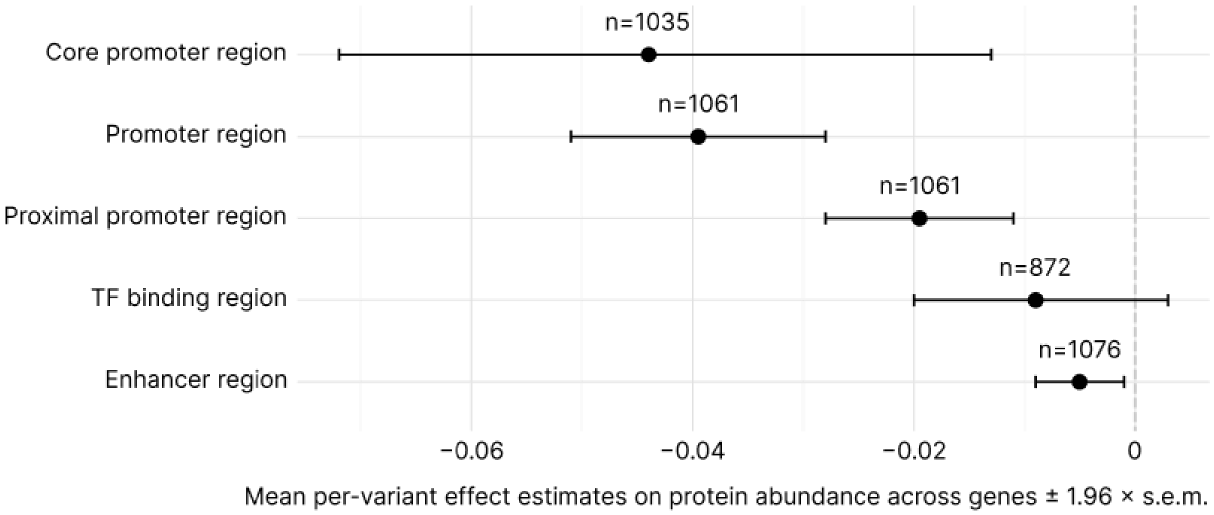


**Supplementary Figure 5. Non-coding variant effects depend on proximity to TSS.**
**a,** Mean across genes of estimated variant effects on protein abundance averaged within each gene, stratified by non-coding variant regulatory context, labeled with sample size (*n*: number of genes) and error bars (1.96 × s.e.m.) across genes. Core promoter was defined as TSS ± 50 bp, Promoter region was defined as ENCODE-annotated PLS or CA-H3K4me3 variants in ±5 kb of the gene, Proximal promoter region was defined as TSS ± 500 bp, TF binding region was defined as ENCODE annotated TF or CA-TF variants in ±5 kb of the gene, and Enhancer region was defined as ENCODE annotated pELS or dELS variants in ±5 kb of the gene.

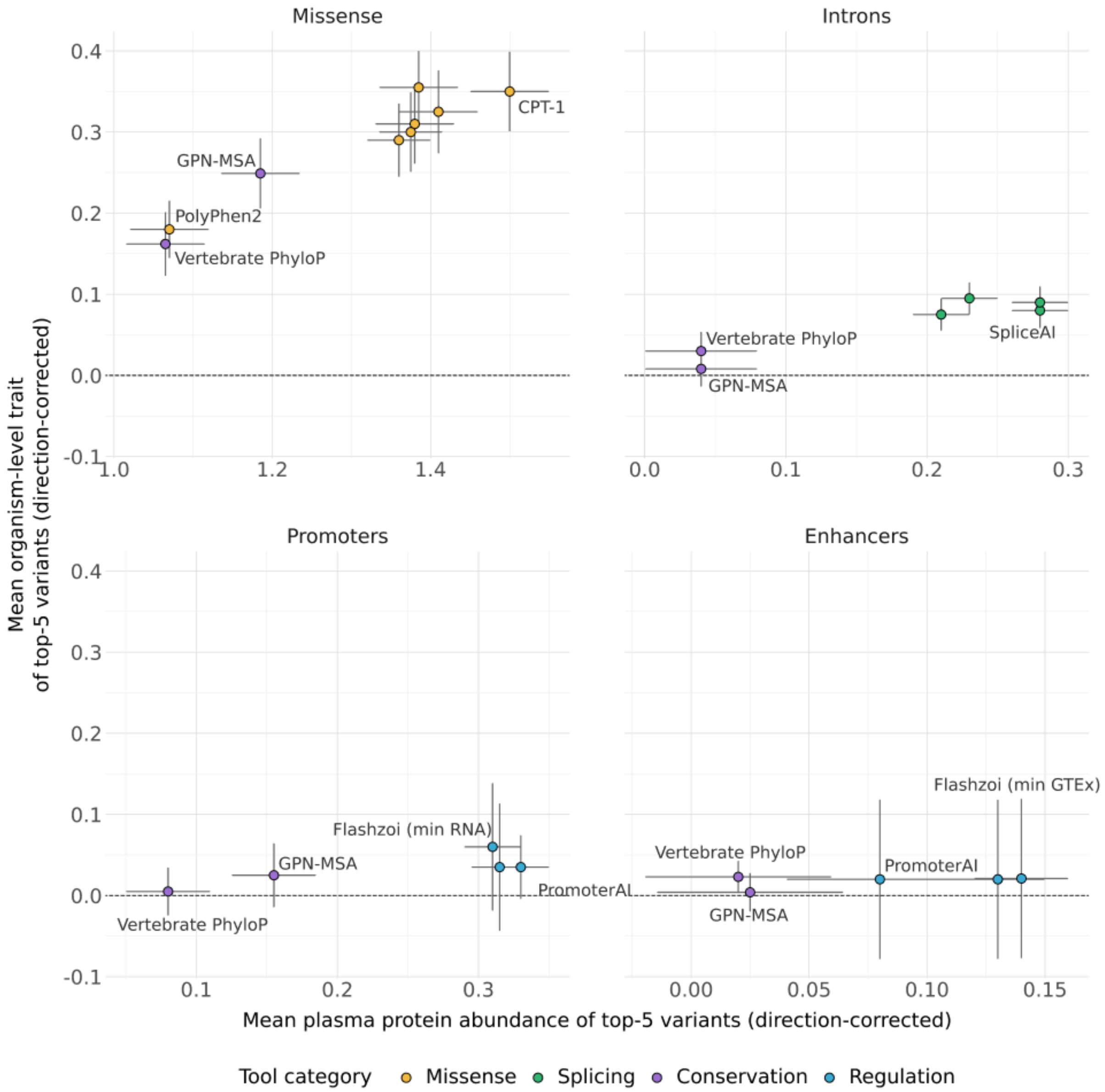


**Supplementary Figure 6: Benchmarking scores using top-5 variants per gene reveals the capacity of variant scores to capture extreme effects.**

Mean across genes of estimated variant effects on quantitative traits (y-axis) versus protein abundance (x-axis) of top-5 variants averaged within each gene according to each scoring method. Facets denote the gene regions from which variants were used. Error bars indicate 1.96 × s.e.m. across genes.

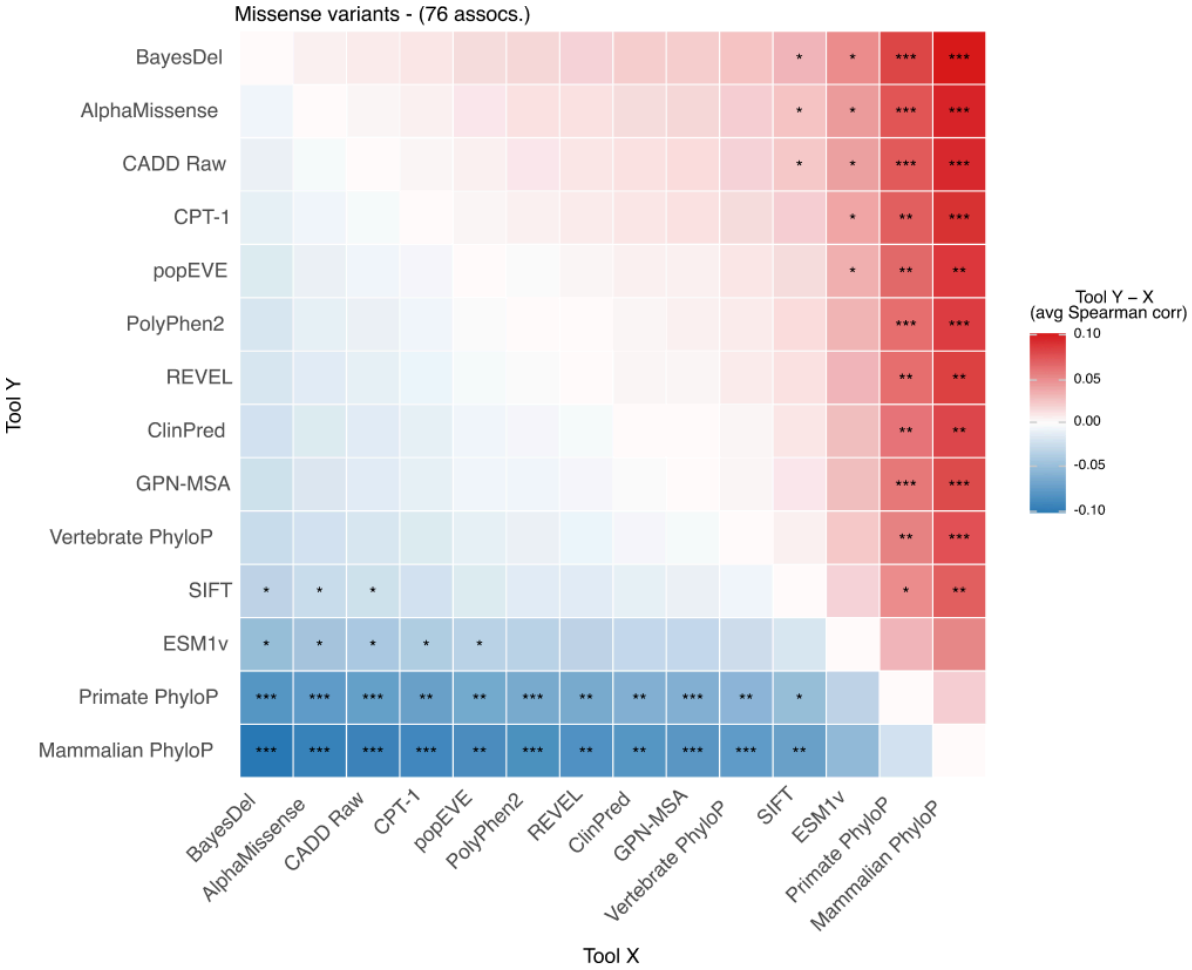


**Supplementary Figure 7: Pairwise comparison of missense variant scores on the All by All META effect estimates.**

Pairwise comparison of Spearman correlations between missense variant scores and All by All variant effect estimates (from the META analysis across genetic ancestries) across 76 gene-trait associations identified on the All of Us cohort. Color indicates the average difference in Spearman correlation (Tool Y − Tool X); significance assessed by the Wilcoxon signed-rank test on the paired per-gene–trait correlations (*** $P < 0.001$, ** $P < 0.01$, * $P < 0.05$).

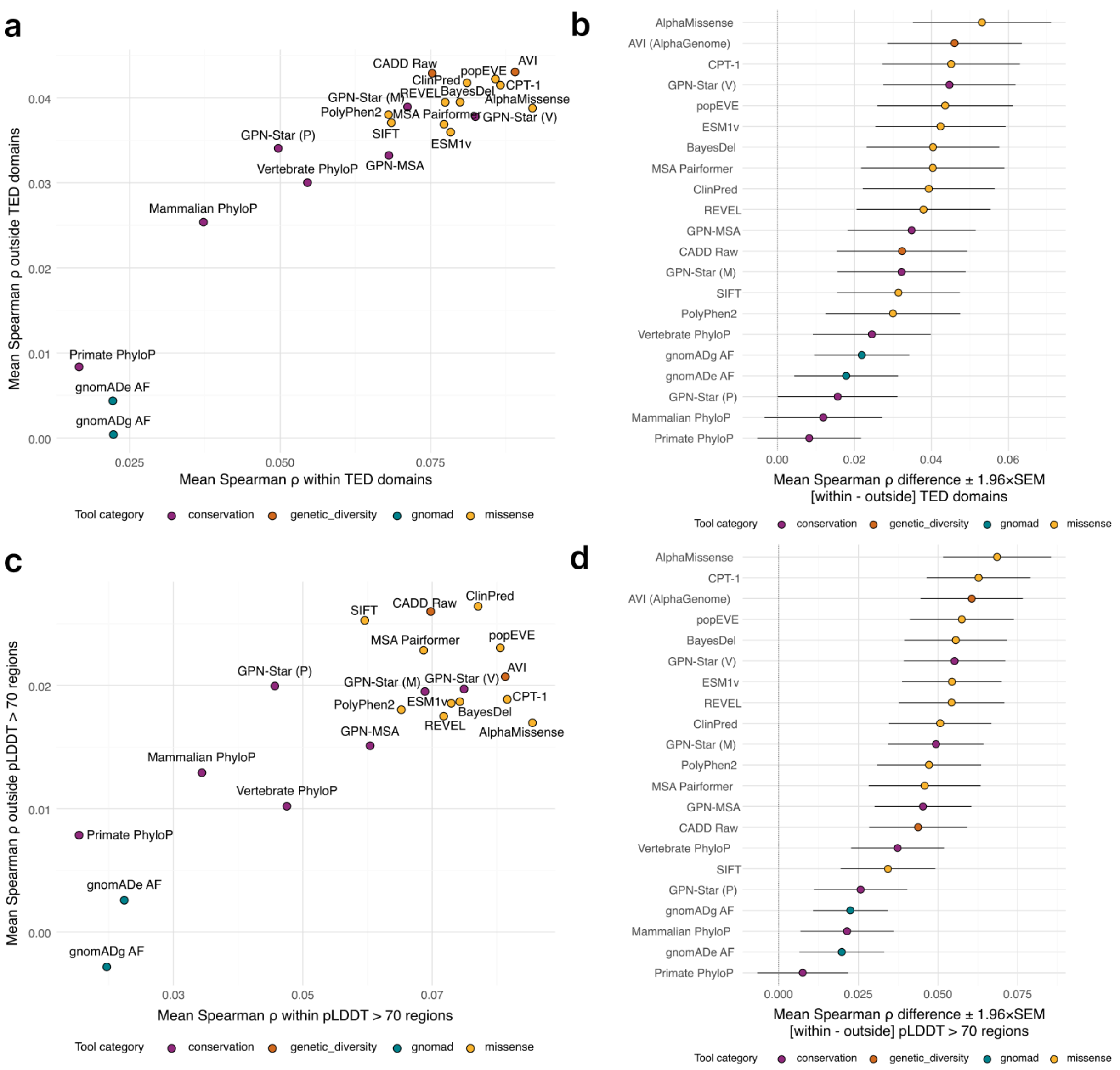


**Supplementary Figure 8: The performance of missense variant scores depends on protein regions.**

**a,** Mean Spearman correlation between each variant score and variant effect estimates on quantitative traits (Genebass) for missense variants located outside TED-defined structured domains (y-axis) versus inside them (x-axis) in the same set of proteins. Each point is a variant score, colored by functional category. **b,** Per-method difference in mean Spearman correlation between missense variants within versus outside TED-defined structured domains [within − outside]. Each point represents a variant-scoring method, colored by functional category; error bars indicate 1.96 × s.e.m. across genes. **c,** Same as in (a), but structured domains are defined as regions with AlphaFold2 pLDDT greater than 70. **d,** Same as in (b), but structured domains are defined as regions with AlphaFold2 pLDDT greater than 70.

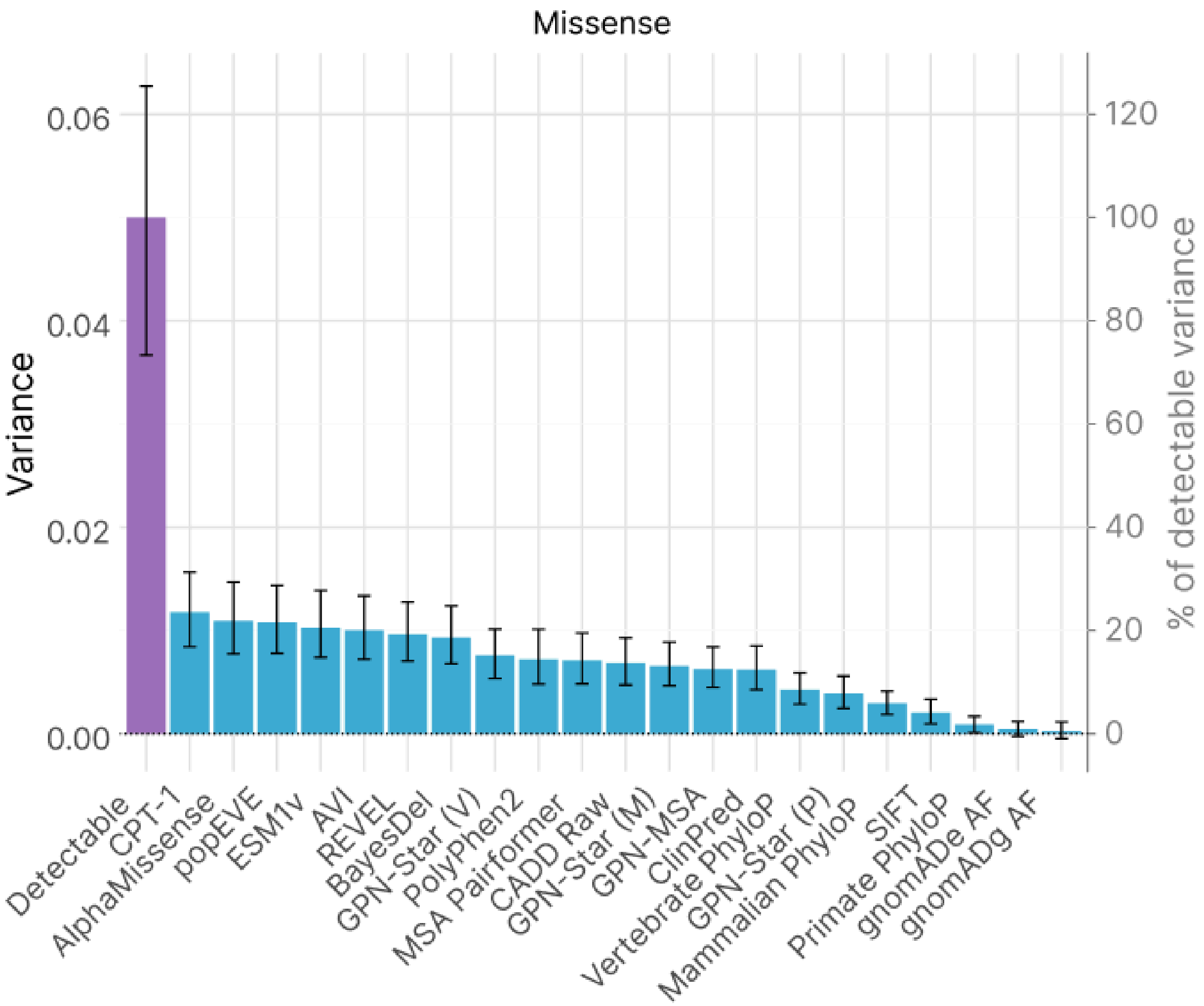


**Supplementary Figure 9: Detectable and captured variance across missense variants.** Variance in variant effect estimates captured by each variant score (primary y-axis) for missense variants. The purple bar indicates the detectable variance, defined as the observed variance minus the estimated sampling-error variance. The percentage of the detectable variance is shown on the secondary y-axis. Error bars indicate 95% confidence intervals from 2,000 gene-level bootstraps.

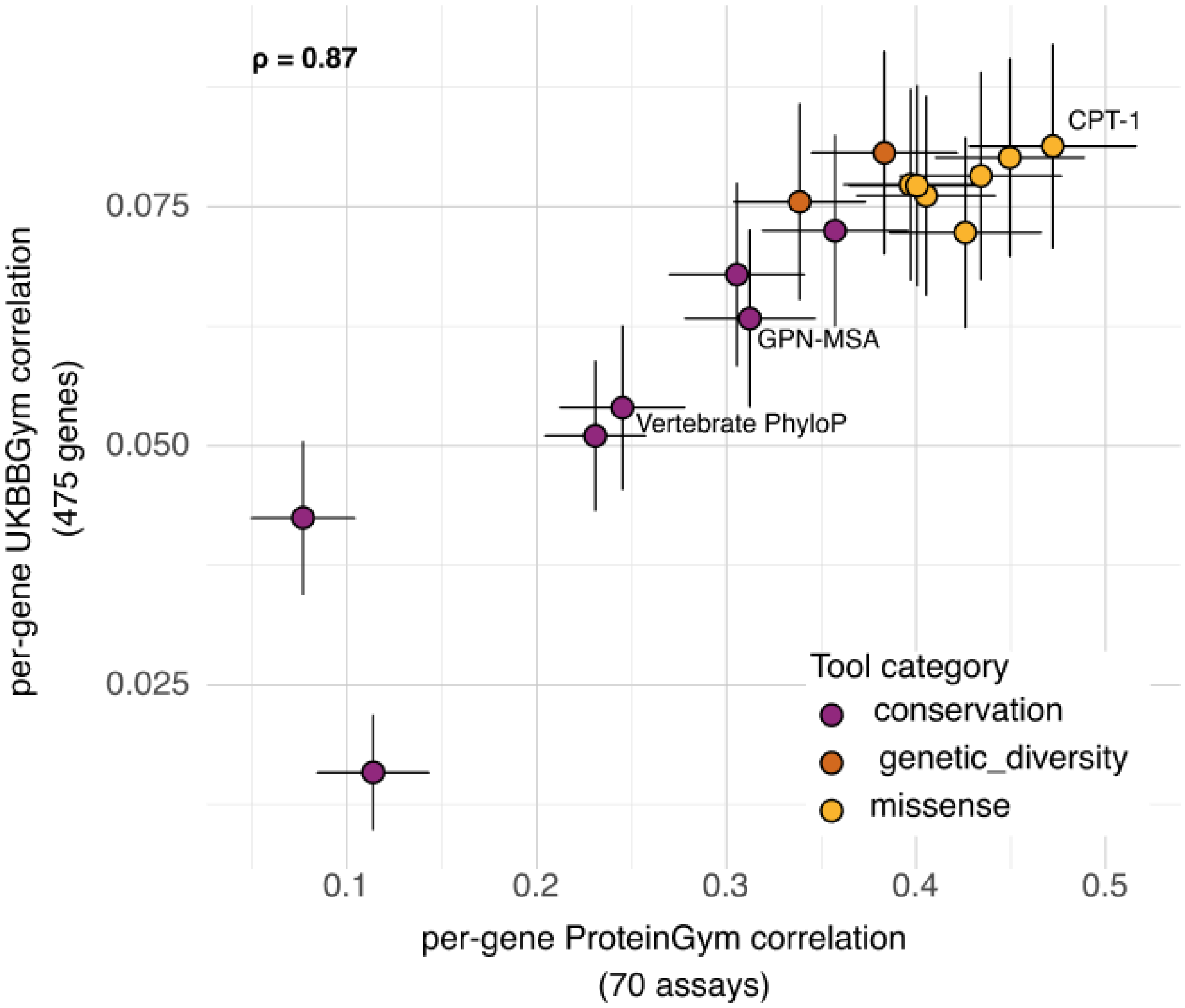


**Supplementary Figure 10: ProteinGym Spearman correlations against UKBBGym Spearman correlations.**

Mean Spearman correlation between each score and variant effect estimates on quantitative traits (Genebass effect estimates; 475 gene-trait pairs) versus mean Spearman correlation of scoring methods on ProteinGym human assay readouts (70 assays with human proteins). Error bars indicate 1.96 × s.e.m. across genes.

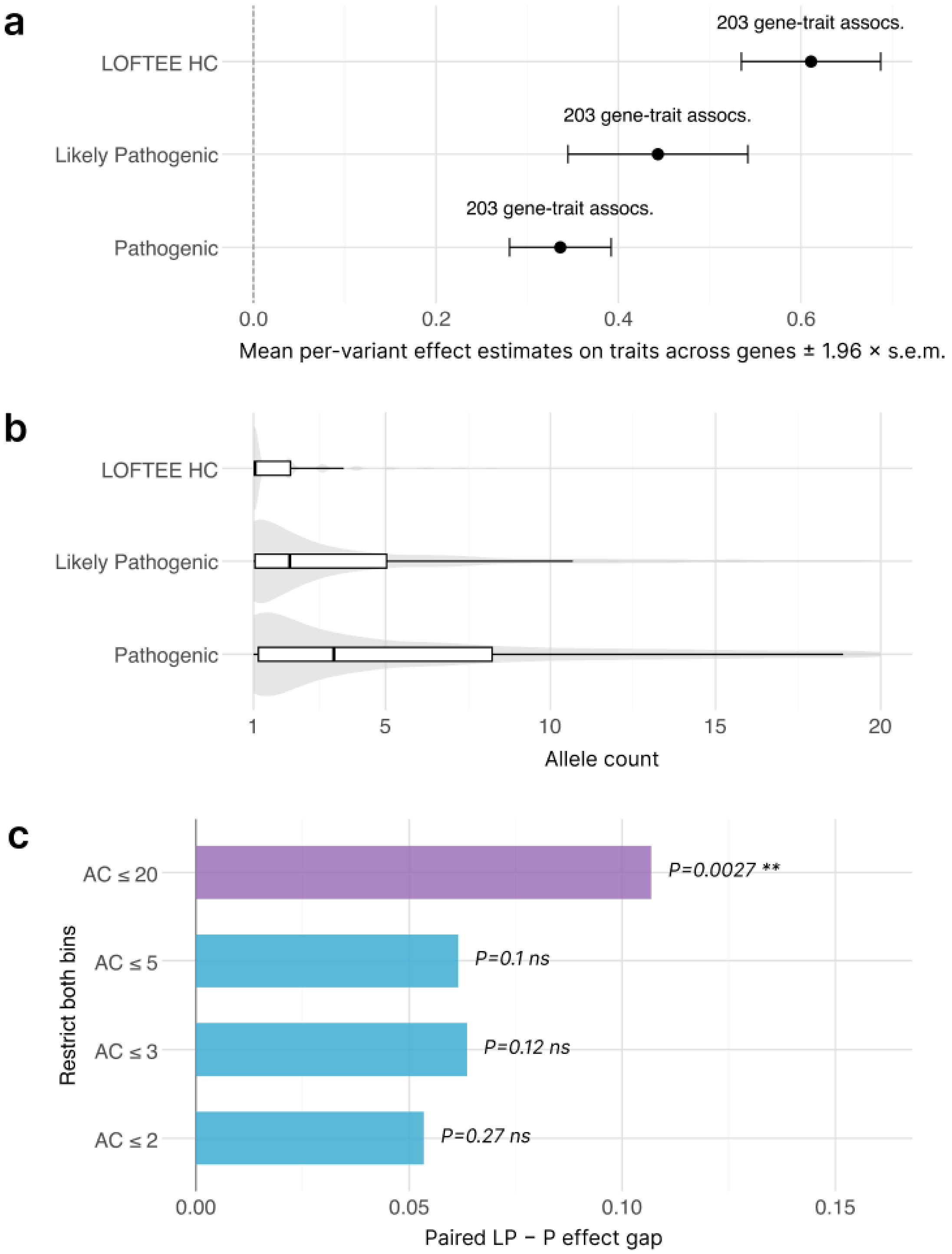


**Supplementary Figure 11. ClinVar likely pathogenic variants do not associate with stronger effects on quantitative traits than pathogenic variants when controlling for allele count.**

**a,** Mean across genes of Genebass variant effect estimates averaged within each gene, for ClinVar Pathogenic, ClinVar Likely Pathogenic, and LOFTEE High Confidence variants. A common set of 203 genes is used across all variant annotations. Error bars indicate 1.96 × s.e.m. **b,** Distribution of Genebass allele counts within each category (box plots over density). **c,** Difference in mean effect estimate between likely pathogenic and pathogenic variants after restricting both categories to a common allele-count ceiling (AC ≤ 20, ≤ 5, ≤ 3, ≤ 2); paired-test p-values are shown (red, significant; blue, non-significant).
