## Supplementary Information for "Phenotypes of ultra-rare variant carriers benchmark variant effect scores"

#### Supplementary Note 1: Relation between the mean phenotype across carriers and summary-statistic variant effect estimates

UKBBGym scores a variant by the average phenotype across its carriers. Published single-variant analyses instead report a regression coefficient (BETA) [40, 66]. This subsection motivates mathematically why the two quantities are approximately the same in the regime we consider them.

For each ultra-rare variant (allele count $\leq 20$), we defined the per-variant phenotype as the mean adjusted phenotype across heterozygous carriers.

For $N$ individuals, let $Y$ be the phenotype vector, $G \in \{0,1\}^N$ the carrier indicator for a given variant (we filtered for heterozygous carriers), and $X$ be the covariate matrix whose first column is all ones (intercept), with the remaining columns age, sex, genetic principal components, and the phenotype-specific polygenic risk score. Let

$$n = \sum_i G_i = G^\top G \tag{1}$$

be the number of carriers and $p = n/N$ the carrier frequency. Let $P_X = X(X^\top X)^{-1}X^\top$ be the projection onto the covariate space and $M_X = I - P_X$ the residual maker, which is symmetric and idempotent, so $M_X^\top M_X = M_X$. Covariate residualization gives $\tilde{Y} = M_X Y$, the adjusted phenotype used throughout UKBBGym.

Our per-variant estimator is the mean of the adjusted phenotype over carriers (the variant phenotype),

$$\hat{\theta} = \frac{1}{n} \sum_{i:\, G_i=1} \tilde{Y}_i = \frac{G^\top M_X Y}{n}. \tag{2}$$

This quantity is to be compared to the published effect estimates.

The published effect estimates is the coefficient $\hat{\beta}$ on $G$ in the fit $Y = Xb + \beta G + \varepsilon$. By the Frisch-Waugh-Lovell theorem [91,92], $\hat{\beta}$ equals the slope of a simple regression of $\tilde{Y}$ on $\tilde{G} = M_X G$,

$$\hat{\beta} = \frac{\tilde{G}^\top \tilde{Y}}{\tilde{G}^\top \tilde{G}}. \tag{3}$$

Expanding $\tilde{G}^\top \tilde{Y} = G^\top M_X^\top M_X Y = G^\top M_X Y$ using $M_X^\top M_X = M_X$ recovers the numerator of $\hat{\theta}$ in (2).

The estimators differ only in their denominators, $G^\top G = n$ for $\hat{\theta}$ and $\tilde{G}^\top \tilde{G} = G^\top M_X G$ for $\hat{\beta}$. Since $P_X$ is also symmetric and idempotent, their ratio is therefore

$$\frac{\hat{\theta}}{\hat{\beta}} = \frac{G^\top M_X G}{G^\top G} = 1 - \frac{\|P_X G\|^2}{n}. \tag{4}$$

Splitting the projection in (4) into the intercept and the remaining centered covariates $\tilde{Z}$ gives $\|P_X G\|^2 = \|P_{\mathbf{1}} G\|^2 + \|P_{\tilde{Z}} G\|^2$. The intercept term evaluates to $p^2 N$, which contributes exactly $p$ after dividing by $n$. The second term contributes $\delta \geq 0$, the fraction of carrier status predictable from age, sex, principal components, and polygenic risk. Carrier status for a rare variant is only weakly dependent of these covariates, so $\delta \approx 0$ and

$$\hat{\theta} \approx (1 - p)\,\hat{\beta}. \tag{5}$$

Intuitively, $\hat{\beta}$ is the difference between the carrier and non-carrier means of $\tilde{Y}$, while $\hat{\theta}$ is the carrier mean alone. For $p \ll 1$, the non-carrier mean is close to the overall mean of $\tilde{Y}$, which is zero by construction. This would not hold when working with common variants.

For UKBBGym variants $p$ is less than $5.1 \times 10^{-5}$ (allele counts less than or equal to 20 and cohort sizes greater than 390,000), so $\theta$ and $\hat{\beta}$ are almost identical for variants considered in UKBBGym.

### Supplementary Note 2: Captured and detectable variance for a score

#### Noise model

Genebass single-variant effect estimates for singleton variants (allele count = 1) contain both genuine between-variant differences in phenotypic effect and sampling error, i.e. phenotypic variability in single-variant carriers that is unrelated to the variant of interest.

For variant $v$ in gene-trait pair $j = (g, t)$, we model the effect estimate $y_{v,j}$ as a true effect $x_{v,j}$ plus sampling error $\epsilon_{v,j}$:

$$y_{v,j} = x_{v,j} + \epsilon_{v,j}. \tag{6}$$

We assume that sampling errors have mean zero, common variance $\sigma_t^2$ within each trait, are uncorrelated across variants, and are uncorrelated with the true effects. Therefore,

$$V_{\text{total}} = V_{\text{true}} + V_{\text{noise}}. \tag{7}$$

Within gene-trait pair $j$ and variant category $c$, we denote these variances by $V_{\text{total},j,c}$ and $V_{\text{true},j,c}$, with

$$V_{\text{noise},t(j)} = \sigma_{t(j)}^2. \tag{8}$$

#### Sampling-error variance

We assumed the sampling error to have a common variance within a trait. We estimated the sampling-error variance $\sigma_t^2$ separately for each trait $t$ from singleton synonymous variants in genes not significantly associated with that trait.

Because these variants are expected to have little systematic phenotypic effect, their observed variance provides an empirical estimate of sampling-error variance at the same allele count and on the same effect-size scale as the analysis variants. No additional calibration or rescaling was applied.

#### Captured variance

For each score $S$, we fitted the following ordinary least-squares regression across variants $v$, separately within each gene-trait pair $j$ and variant category $c$:

$$y_{v,j} = \alpha_{c,j} + \beta_{c,j} S_{v,j} + \eta_{v,j}. \tag{9}$$

This gave residuals $\hat{\eta}_{v,j}$.

The explained sum of squares is

$$SS_{\text{explained},j,c} := SS_{\text{total},j,c} - SS_{\text{resid},j,c}, \tag{10}$$

that is,

$$SS_{\text{explained},j,c} = \sum_{v \in c} (y_{v,j} - \bar{y}_{j,c})^2 - \sum_{v \in c} \hat{\eta}_{v,j}^2. \tag{11}$$

We defined the **captured variance** by pooling the sampling-error-corrected explained sum of squares across gene-trait pairs and dividing by the corresponding pooled degrees of freedom:

$$\hat{V}_{\text{captured,c}} := \frac{\sum_j \left[ SS_{\text{explained},j,c} - \sigma_{t(j)}^2 \right]}{\sum_j (n_{j,c} - 1)}. \tag{12}$$

#### Detectable variance

We defined the **detectable variance** as the expected captured variance (12) for a **perfect linear score**, i.e. a score for which, within every gene-trait pair and category, there exist coefficients $a_{c,j}$ and $b_{c,j}$ such that

$$x_{v,j} = a_{c,j} + b_{c,j} S_{v,j}. \tag{13}$$

For a perfect linear score, the true effects satisfy the linear regression model exactly, so the regression error in (6) is given by the sampling error $\epsilon_{v,j}$. Therefore,

$$\mathbb{E}[SS_{\text{resid},j,c}] = (n_{j,c} - 2)\sigma^2_{t(j)}. \tag{14}$$

Moreover,

$$\begin{aligned} \mathbb{E}[SS_{\text{total},j,c}] &= (n_{j,c} - 1)V_{\text{total},j,c} \\ &= (n_{j,c} - 1)\left[V_{\text{true},j,c} + \sigma^2_{t(j)}\right]. \end{aligned} \tag{15}$$

Hence,

$$\begin{aligned} \mathbb{E}[SS_{\text{explained},j,c}] &= (n_{j,c} - 1)\left[V_{\text{true},j,c} + \sigma^2_{t(j)}\right] - (n_{j,c} - 2)\sigma^2_{t(j)} \\ &= (n_{j,c} - 1)V_{\text{true},j,c} + \sigma^2_{t(j)}. \end{aligned} \tag{16}$$

Subtracting the expected sampling-error contribution $\sigma^2_{t(j)}$ from (16) gives

$$\mathbb{E}\left[SS_{\text{explained},j,c} - \sigma^2_{t(j)}\right] = (n_{j,c} - 1)V_{\text{true},j,c}. \tag{17}$$

Altogether,

$$\begin{aligned} V_{\text{detectable},c} &:= \mathbb{E}\left[\hat{V}_{\text{captured},c} \mid S \text{ is a perfect linear score}\right] \\ &= \frac{1}{W_c}\sum_j (n_{j,c} - 1)V_{\text{true},j,c}, \end{aligned} \tag{18}$$

where

$$W_c = \sum_j (n_{j,c} - 1). \tag{19}$$

Thus, the detectable variance is the weighted average of the true between-variant effect variance across gene-trait pairs, with the degrees of freedom $n_{j,c} - 1$ as weights.

An empirical estimator of the detectable variance (18) is obtained from the observed total sum of squares after subtracting the expected sampling-error contribution:

$$\hat{V}_{\text{detectable},c} = \frac{\sum_j \left[SS_{\text{total},j,c} - (n_{j,c} - 1)\sigma^2_{t(j)}\right]}{\sum_j (n_{j,c} - 1)}. \tag{20}$$

Equation (20) estimates the detectable variance and provides the empirical noise ceiling against which the captured variance (12) of each score is compared.

#### Orthogonal check on the reported standard

As a diagnostic, we compared the empirical trait-specific noise variance with the mean squared Genebass standard error for the same singleton synonymous variants. Their ratio had a median of 1.04 across 100 traits (10th-90th percentile, 0.96-1.12), indicating close agreement between the empirical noise estimate and the reported standard errors. This comparison was diagnostic only and was not used in downstream analyses.

#### Supplementary Tables

##### Supplementary Table 1.

Significant gene-quantitative trait associations used in UKBBGym. These were found through LOFTEE-HC burden tests on the same data.

##### Supplementary Table 2.

Genes in which LOFTEE-HC significantly reduced adjusted Olink plasma protein abundance.

##### Supplementary Table 3.

Variant scores benchmarked and genomic regions annotations used within UKBBGym. The table contains the sources of the scores and annotations, and the neutral values used as the method's neutral effect score.

##### Supplementary Table 4.

Significant gene-quantitative trait associations used in the All of Us implementation of UKBBGym. These were found through LOFTEE-HC burden tests on the same data in the All by All study.
